# Type I interferon governs immunometabolic checkpoints that coordinate inflammation during *Staphylococcal* infection

**DOI:** 10.1101/2024.01.10.575104

**Authors:** Mack B. Reynolds, Benjamin Klein, Michael J. McFadden, Norah K. Judge, Hannah E. Navarrete, Britton C Michmerhuizen, Dominik Awad, Tracey L. Schultz, Paul W. Harms, Li Zhang, Teresa R. O’Meara, Jonathan Z. Sexton, Costas A. Lyssiotis, J. Michelle Kahlenberg, Mary X. O’Riordan

**Author notes:** Correspondence: MaryX. O’Riordan, PhD, Department of Microbiology and Immunology, University of Michigan Medical School, Ann Arbor, Michigan, USA.

## Abstract

Fine-tuned inflammation during infection enables pathogen clearance while minimizing host damage. Inflammation regulation depends on macrophage metabolic plasticity, yet coordination of metabolic and inflammatory programs is underappreciated. Here we show that type I interferon (IFN) temporally guides metabolic control of inflammation during methicillin-resistant Staphylococcus aureus (MRSA) infection. In macrophages, staggered Toll-like receptor and type I IFN signaling permitted a transient respiratory burst followed by inducible nitric oxide synthase (iNOS)-mediated OXPHOS disruption. OXPHOS disruption promoted type I IFN, suppressing other pro-inflammatory cytokines, notably IL-β. iNOS expression peaked at 24h post-infection, followed by lactate-driven iNOS repression via histone lactylation. In contrast, type I IFN pre-conditioning sustained infection-induced iNOS expression and amplified type I IFN. Cutaneous MRSA infection in mice constitutively expressing epidermal type I IFN led to elevated iNOS levels, impaired wound healing, vasculopathy, and lung infection. Thus, kinetically regulated type I IFN signaling coordinates immunometabolic checkpoints that control infection-induced inflammation.

**Graphical abstract:** 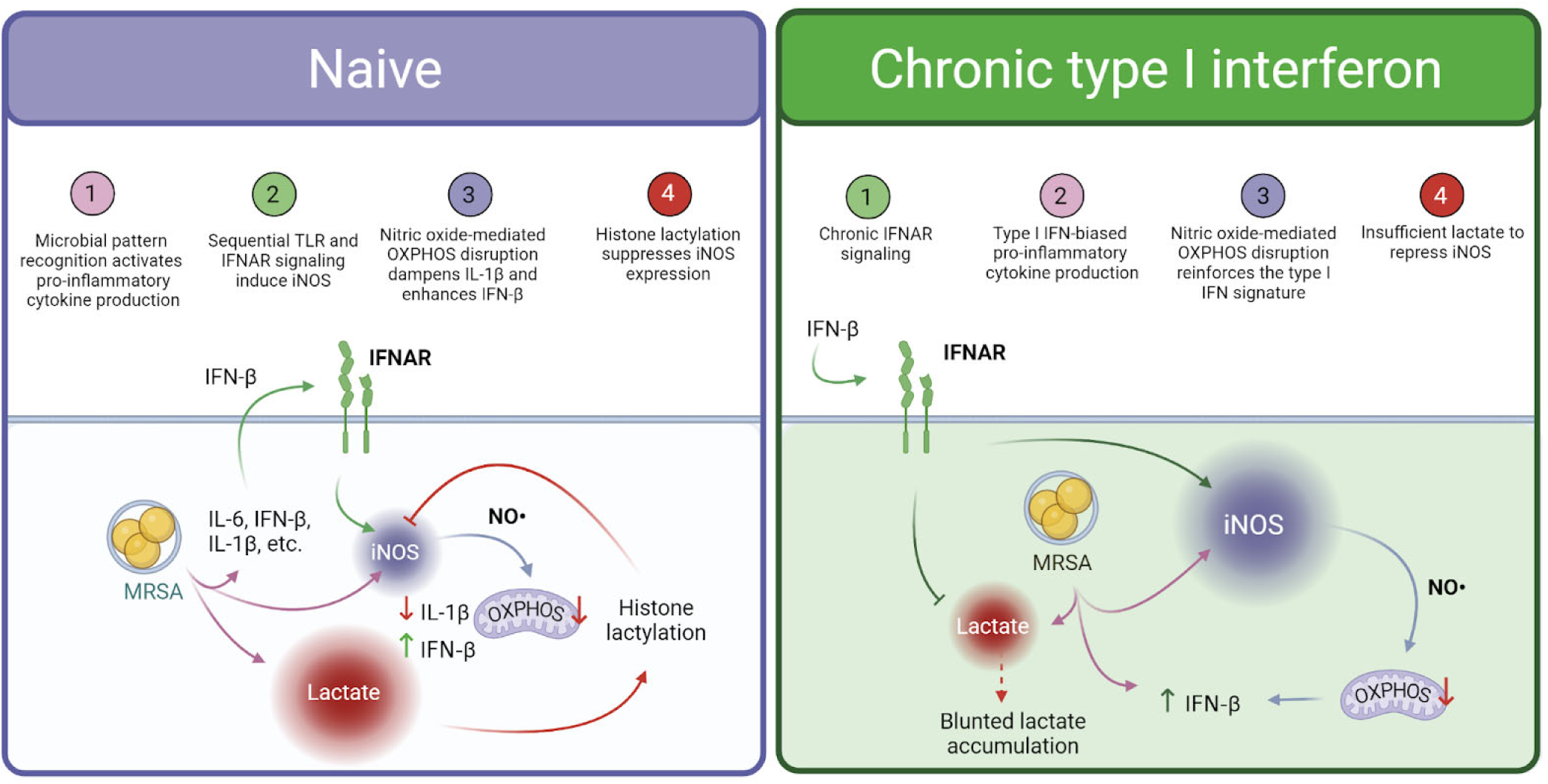

## Introduction

Metabolic remodeling in immune cells guides initiation and resolution of inflammation, balancing the benefits of inflammation with the consequences of immune-mediated damage^1–4^. As innate immune cells tasked with regulating inflammation, macrophages exhibit remarkable metabolic plasticity in health and disease^4–13^. During bacterial infection, macrophages adapt their metabolism to regulate bioenergetics, signaling, inflammatory gene expression, and antimicrobial function. Productive host defense depends on fine-tuning these potentially competing metabolic outputs. A common paradigm observed in macrophages activated by bacterial infection is a switch from oxidative phosphorylation (OXPHOS) to aerobic glycolysis-driven adenosine triphosphate (ATP) production^14–20^. While a common interpretation is that this metabolic switch acts to rapidly generate energy, this profound shift may reflect more complex metabolic changes with pleiotropic functional consequences^21–24^. Thus, a comprehensive analysis of the sequence and regulation of metabolic reprogramming that occurs in macrophages during infection will lend valuable insight into the metabolic coordination of inflammation.

The innate immune response to bacterial pathogens is governed by highly conserved pattern recognition receptors, such as the Toll-like receptors (TLR)^25^. Additionally, most microbial infections trigger production of type I IFN (IFN) through TRIF or nucleic acid sensing^26^. However, type I IFN plays a complex role in antibacterial immunity, as IFN can either enhance or decrease susceptibility to various bacterial pathogens^26–29^. Recent work highlights a critical role of IFN in controlling macrophage metabolism during bacterial infection^15^. Furthermore, chronic overproduction of type I IFN, such as in the autoimmune disease systemic lupus erythematosus, or in viral co-infections, can enhance susceptibility to bacterial infection and associated inflammatory disease^30–33^. Thus, IFN responses must be well-calibrated during bacterial infection to facilitate host defense without excess host damage, a general principle known as the damage-response framework^34^. One hypothesis that could explain the complex role of type I IFN in bacterial infection is that magnitude or timing of type I IFN signaling distinctly shapes host metabolism to promote or impair antibacterial immunity.

Infection with the bacterial pathogen *Staphylococcus aureus* elicits a rapid and robust innate immune response’ characterized by enhancement of glycolytic metabolism and itaconate-dependent TCA cycle remodeling^17 35–38^. During S. *aureus* skin infection, dermal macrophages cycle between a GM-CSF and HIF-1a-driven homeostatic metabolic program and an itaconate-dominated pro-inflammatory metabolic program^39^. These changes in host metabolism facilitate antibacterial immunity, inflammation, immune memory, and inflammation resolution during S. *aureus* infection. At the same time, common comorbidities can impair *anti-Staphylococcal* immunity, including viral co-infection, autoimmune disease, and metabolic disorders such type II diabetes^30,40^ ^42^. Common among these disease states, dysregulation of macrophage metabolism may be an underlying condition that impairs immunity to S. *aureus* infection.

Here, we demonstrate that S. *aureus* infection in macrophages elicits a series of kinetically staggered metabolic programs that guide pro-inflammatory cytokine responses. Early toll-like receptor (TLR) signaling drives an energetic burst of OXPHOS and aerobic glycolysis. Later, dual TLR and IFN signaling triggers a sharp reduction in OXPHOS via nitric-oxide-mediated disruption of the electron transport chain (ETC) with maintenance of high levels of aerobic glycolysis. We demonstrate that these sequential metabolic shifts support macrophage inflammatory programming during S. *aureus* infection, including regulation of type I IFN and IL-β. Further, while extensive metabolic remodeling occurs in S. at/reus-infected macrophages, our data support that two metabolites, nitric oxide and lactate, serve as immunometabolic checkpoints for macrophage metabolic reprogramming and pro-inflammatory cytokine responses. Finally, we establish type I IFN as a key pacesetter for these immunometabolic checkpoints during S. *aureus* infection. Taken together, our findings lend new insight into the cross-talk between immune signaling and metabolic programming in macrophages that enables their immunoregulatory role in inflammation and host defense.

## Results

### MRSA infection dampens macrophage OXPHOS through electron transport chain disruption

Bacterial infection provokes profound metabolic remodeling in macrophages that may be influenced by differential pattern recognition receptor engagement, inflammatory signaling, and bacteria-specific determinants^14,43–45^. To define global energetic changes driven by methicillin-resistant *Staphylococcus aureus* (MRSA) infection, we infected murine bone marrow-derived macrophages (BMDM) with MRSA for 24 h and performed a Seahorse extracellular flux (XF) assay to measure rates of oxygen consumption and extracellular acidification as proxies for oxidative phosphorylation (OXPHOS) and glycolysis, respectively. By 24 h, MRSA infection shut down mitochondrial respiration **(Fig 1** and **Fig S1A-C)** without a significant decrease in macrophage survival or change in levels of the mitochondrial outer membrane protein TOM20 **(Fig S1D-F).** We hypothesized that MRSA infection could directly disrupt the electron transport chain (ETC). To test this hypothesis, we measured the abundance of the respiratory complexes (RC), the functional units of the ETC, using blue native-polyacrylamide gel electrophoresis (BN-PAGE) and immunoblot with a panel of antibodies targeting subunits from each complex **(Fig 1B-C).** MRSA infection triggered loss of Complex I, Complex II, Complex IV, and super-respiratory complexes (SRC) but did not affect the Complex III dimer or Complex V. To test the 5 effect of MRSA infection on macrophage mitochondrial polarization, we used flow cytometry to track staining with a mitochondrial membrane potential-sensitive dye (MitoTracker Deep Red) and a membrane potential-independent dye (MitoTracker Green) **(Fig 1D and Fig S1G),** which allows differentiation between changes in mitochondrial polarization and mass. Under basal conditions, membrane potential scaled linearly with mitochondrial content. However, MRSA infection triggered mitochondrial hyperpolarization, indicating ETC disruption might have been preceded by a period of elevated OXPHOS^46^. Our data point to destabilization of the RCs as the likely route of OXPHOS disruption in MRSA-infected macrophages.

**Figure 1.**
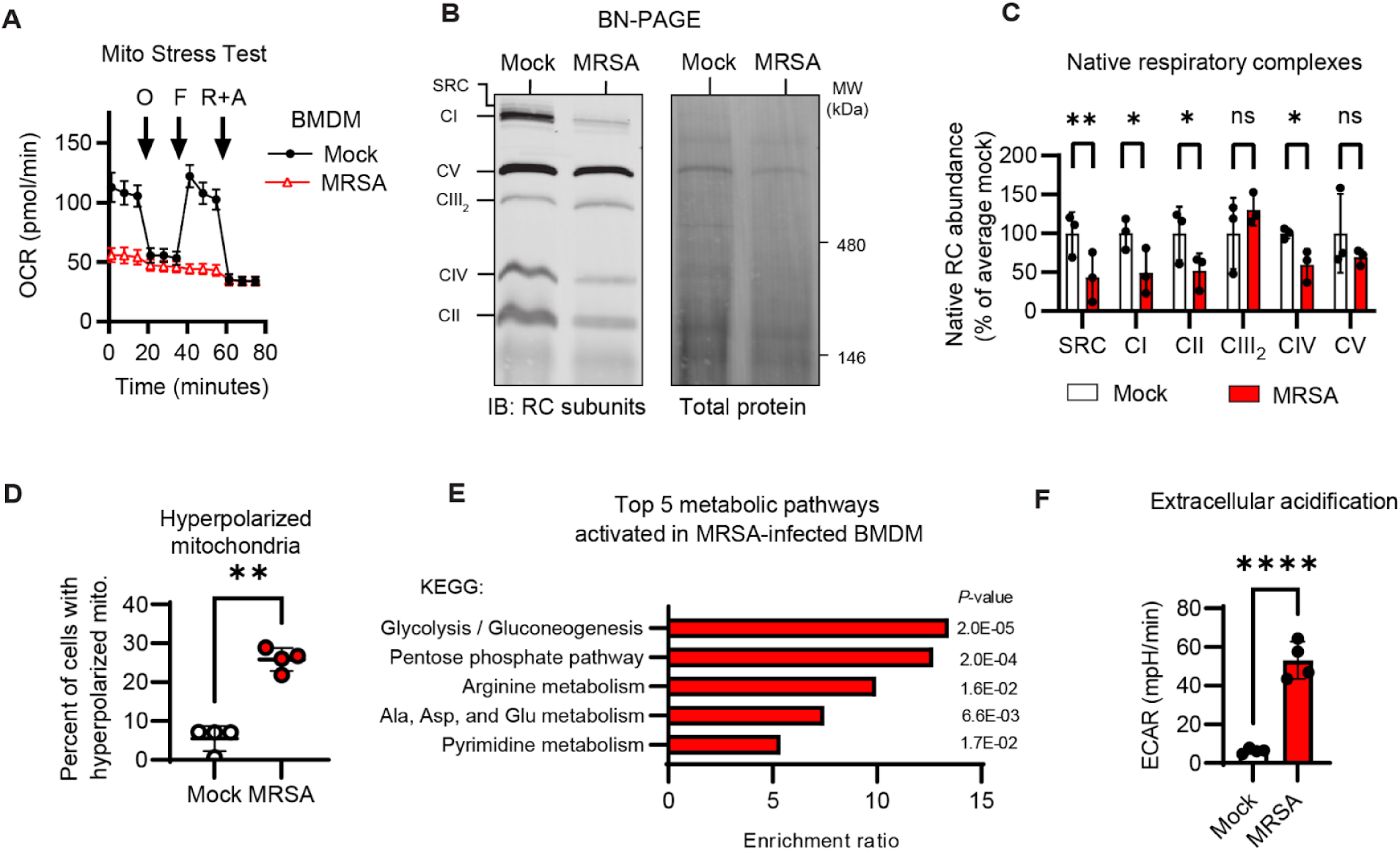
MRSA infection disrupts the electron transport chain at multiple points and stimulates robust glycolysis in macrophages. **A.** Seahorse extracellular flux (XF) analysis of the rate of oxygen consumption (OCR) in 24 h MRSA-infected (USA300; MOI 20) BMDM using the Mito Stress Test assay, with additions of oligomycin (O), carbonyl cyanide p-trifluoromethoxyphenylhydrazone (FCCP), rotenone (R), and antimycin A (A) at the indicated time points. B. Blue native (BN)-PAGE and immunoblot analysis of native respiratory complexes (RC) Complex l-V (Cl-V) and super-respiratory complexes (SRC) paired to total protein stain (Coomassie G-250) in mock or 24 h MRSA-infected BMDM. **C.** Quantification RC abundance relative to total protein as a percent of the average mock samples between experiments. D. Flow cytometric analysis of mitochondrial polarization following dual staining with a membrane potential sensitive mitochondrial dye (MitoTracker Deep Red; MTDR) and a membrane potential insensitive mitochondrial dye (MitoTracker Green; MTG) in mock and 24 h MRSA-infected BMDM. Quantification of the percent of MTDR^High^ (hyperpolarized) cells. E. LC-MS-based targeted metabolomics analysis of 230 metabolites in mock or 24 h MRSA-infected BMDM with Metaboanalyst 5.0 pathway enrichment analysis using KEGG annotation of the top 5 significantly changed metabolic pathways between mock and MRSA-infected BMDM. **F.** Seahorse XF analysis of the rate of extracellular acidification (ECAR) in mock or 24 h MRSA-infected BMDM. Graphs represent the mean of n > 3 biological replicates with SD error bars. P-values were calculated using an unpaired t test **(D** and F) or two-way analysis of variance (ANOVA) with Sidak’s post test (C). *P < 0.05; **P < 0.01; ***‘P < 0.0001.

To define metabolite changes elicited by MRSA infection, we infected BMDM with MRSA for 24 h and extracted intracellular metabolites for liquid chromatography-mass spectrometry (LC-MS)-based targeted metabolomics of 230 well-characterized mammalian metabolites **(Fig S1H-L** and **File S1).** Glucose metabolism, the pentose phosphate pathway, and arginine metabolism were the most significantly altered metabolic pathways in MRSA-infected macrophages **(Fig 1E).** Metabolites characteristic of macrophage inflammatory activation were robustly stimulated, including itaconate and L-citrulline, products of the infection-induced enzymes immune responsive gene 1 (IRG1) and inducible nitric oxide synthase (iNOS) respectively **(Fig S1L)**^47 48^. Furthermore, we noted increased intracellular lactate, despite high levels of extracellular acidification resulting from lactate release **(Fig 1F** and **Fig S1L).** MRSA infection significantly increased succinate, likely due to loss of Complex II (succinate dehydrogenase) or Complex II inhibition by itaconate. Notably, macrophage metabolic remodeling sustained high ATP levels, despite OXPHOS shutdown **(Fig S1L),** indicating that macrophages are fully adapted energetically to this profound alteration in the route of ATP production. Collectively, these data reinforce macrophage OXPHOS shutdown and the concomitant glycolytic shift as characteristic of MRSA infection.

### Staggered TLR and IFNAR signaling coordinate a transient burst of OXPHOS followed by OXPHOS shutdown during MRSA infection

Bacterial pattern recognition through toll-like receptors (TLR) and cytokine signaling through JAK-STAT pathways are classical signatures of macrophage inflammatory activation^49^. We hypothesized that dampened OXPHOS in MRSA-infected macrophages occurs through engagement of one or both pathways^15^. We first measured individual contributions of TLR or IFNAR signaling to ETC disruption by 7’ stimulating BMDM with the TLR2 agonist Pam3CSK4 (P3CSK4) with or without IFN-β and performing a Seahorse XF assay **(Fig 2A-B).** TLR2 activation alone resulted in a robust increase in basal respiration, while IFN-β alone did not affect respiration. Co-stimulation with both TLR2 and IFNAR agonists led to a substantial drop in basal respiration, phenocopying MRSA infection at 24 h. In parallel, we tested the relative contributions of TLR2 and/or IFN signaling to mitochondrial hyperpolarization and found that mitochondrial hyperpolarization only occurred upon combined TLR2/IFN signaling **(Fig S2A-B).** Additionally, we found that *Tlr2/4/9-l-* immortalized BMDM (iBMDM) did not develop mitochondrial hyperpolarization during MRSA infection or combined TLR2/IFN signaling, indicating that TLR signaling is required for perturbation of the ETC. Finally, we tested the effect of TLR and IFN signaling on assembly and abundance of the RCs that constitute the ETC using BN-PAGE and immunoblot with a panel of RC subunit antibodies **(Fig 2C-D).** As with mitochondrial membrane potential hyperpolarization, both TLR and IFN signaling were needed to trigger RC disassembly. Finally, we tested whether RC disassembly occurred through loss of individual RC subunits using denaturing SDS-PAGE and immunoblot against representative subunits of each RC. We determined that subunits of Complex I, Complex II, and Complex IV were decreased at the protein level by either MRSA infection or combined TLR2/IFN signaling **(Fig S2C-D).**

**Figure 2.**
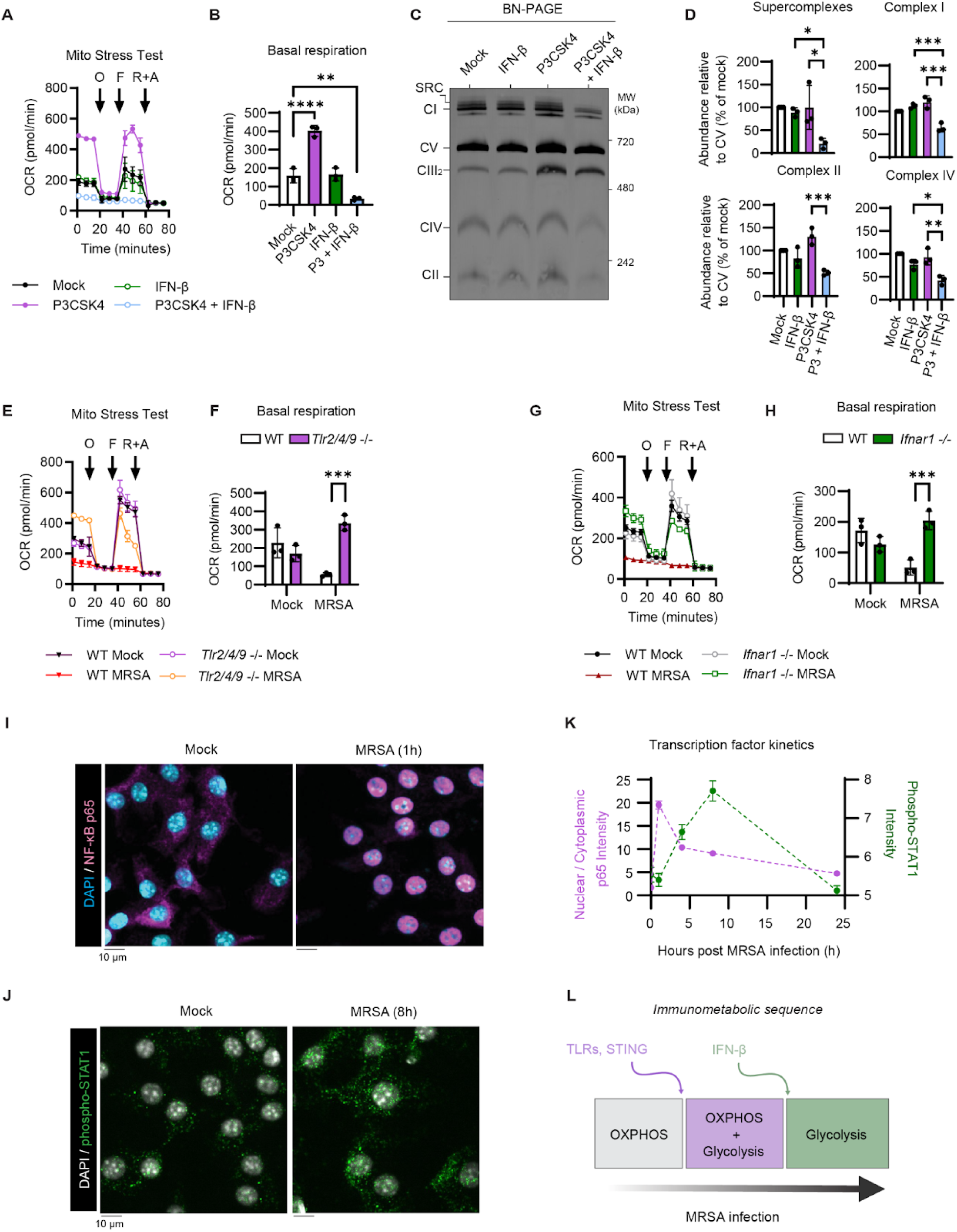
Kinetically staggered TLR and IFNAR signaling set the pace of macrophage metabolism during MRSA infection. **A.** Seahorse XF analysis of BMDM treated for 24 h with TLR2 agonist Pam3CSK4 (P3CSK4; 2 pg/mL) and/or IFN-β (400U/mL) using the Mito Stress Test assay, with additions of oligomycin (O), carbonyl cyanide p-trifluoromethoxyphenylhydrazone (FCCP), rotenone (R), and antimycin A (A) at the indicated time points. B. Quantification of basal respiration in BMDM stimulated for 24 h with or without P3CSK4 and IFN-β. **C.** BN-PAGE and immunoblot analysis of native RC from BMDM treated with P3CSK4 and/or IFN-β for 24 h. **D.** Native RC abundance relative to CV, which does not change upon stimulation, normalized to the paired mock sample. E. Seahorse Mito Stress Test analysis of mock or 24 h MRSA-infected TLR2, 4, 9 triple KO immortalized BMDM *(Tlr2/4/9* -/- iBMDM). F. Basal respiration of mock or 24 h MRSA-infected WT or *Tlr2/4/9* -/- iBMDM. G. Seahorse Mito Stress Test analysis of mock or 24 h MRSA-infected Interferon alpha/beta receptor 1 KO *(Ifnarl* -/-) IBMDM H. Basal respiration quantification in mock or 24 h MRSA-infected WT or *Ifnarl -/-* iBMDM. Representative confocal micrographs from high-content imaging of NF-κB p65 (I) and phospho-STAT1 (J) immunofluorescence staining of BMDM infected with MRSA (MOI 20) for the indicated time. K. Automated analysis of the average nuclear:cytoplasmic ratio of NF-κB p65 and intensity of phospho-STAT1 per cell in CellProfiler. L. Graphical representation of the “immunometabolic sequence,” highlighting signaling pathways and metabolic changes which occur within the first 24 h of MRSA infection, created with Biorender.com. Graphs represent the mean of n > 3 biological replicates with SD error bars, except **Fig 2K** which represents the mean and 95% confidence interval of >492 cells per condition pooled across 3 biological replicates. P-values were calculated using a one-way ANOVA with Tukey’s post test (B and D) or two-way ANOVA with Sidak’s post test **(F** and **H).** *P < 0.05; **P < 0.01; ***P < 0.001; ****P < 0.0001.

After determining that TLR and IFN signaling were sufficient to trigger ETC disruption, we tested if these pathways were necessary for ETC disruption during MRSA infection. We infected *Tlr2/4/9* -/- and WT iBMDM with MRSA for 24 h and analyzed OXPHOS using a Seahorse XF assay **(Fig 2E-F).** TLR signaling was required for MRSA-induced OXPHOS suppression. Further, we found that MRSA-induced loss of RC subunits SDHB, NDUFB8, and MTCO1 depended on TLR signaling **(Fig S2E-F).** Next, we infected *Ifnarl -I-* and WT iBMDM with MRSA for 24 h and performed a Seahorse XF assay **(Fig 2G-H).** Like TLR signaling, IFNAR signaling was required for MRSA-induced OXPHOS disruption and loss of respiratory complex subunits SDHB, NDUFB8, or MTCO1 **(Fig S2G-H).** Taken together, our data support that combined TLR and IFNAR signaling is required for ETC disruption during MRSA infection.

During a physiological MRSA infection, the TLR and IFNAR pathways would be activated sequentially, with TLR2 engaged during initial bacterial:host contact, followed by production of type I IFN that stimulates IFNAR. Therefore, we hypothesized that kinetic staggering of these unique signals would establish an inflammatory checkpoint for this profound alteration of mitochondrial metabolism. Using immunofluorescence and automated confocal microscopy, we tracked kinetics of downstream TLR and IFNAR signaling, through NF-κB nuclear translocation and STAT1 phosphorylation, respectively **(Fig 2I-K** and **Fig S2I-J).** We observed a kinetically staggered response, where NF-κB nuclear translocation was elevated within the first hour and STAT1 phosphorylation was elevated at 8 h post-infection. Notably, maximal STAT1 phosphorylation coincided with the highest detected levels of MRSA-induced secreted IFN-β **(Fig S2K).** Collectively, these observations suggest the model of an immunometabolic circuit **(Fig. 2L),** whereby sequential inflammatory signals, first TLR, followed by IFNAR signaling, tune OXPHOS activity.

### IFN signaling drives iNOS expression during MRSA infection to fuel nitric oxide-mediated OXPHOS disruption

As our data pointed to IFNAR as a critical regulator of MRSA-induced macrophage bioenergetics, we next investigated mechanisms by which IFNAR signaling regulates MRSA-induced metabolic remodeling in macrophages. To this end, we performed targeted metabolomics of MRSA-infected WT and *Ifnarl-/-* macrophages **(Fig S3A-C** and **File S2).** Globally, we found that IFNAR was critical for the regulation of metabolites related to arginine and glucose metabolism. **(Fig 3A).** Additionally, prior work demonstrated that OXPHOS disruption in classically activated M1 macrophages occurs through iNOS expression and nitric oxide (NO*)-mediated respiratory complex disassembly^50 51^. Indeed, we observed that IFNAR was a critical regulator of MRSA-induced L-citrulline production and L-arginine consumption **(Fig 3B),** a metabolic signature characteristic of iNOS activity^48^. To specifically test the role of IFNAR on iNOS activity, we measured production of nitrite, a byproduct of NO*, with a Griess assay in MRSA-infected WT and *Ifnarl* -/- iBMDM treated with or without an iNOS inhibitor, L-NIL **(Fig 3C).** L-NIL treatment or IFNAR1 deletion prevented nitrite production. Thus, we conclude that IFNAR controls MRSA-induced arginine metabolism through iNOS. To test the hypothesis that iNOS regulates OXPHOS in MRSA-infected macrophages, we first determined the regulators of iNOS expression during MRSA infection. We stimulated macrophages with TLR2 agonist P3CSK4 with or without IFN-β or the type II IFN, IFN-γ **(Fig S3D-E).** Combined signaling through TLR2 and IFNAR was sufficient to induce iNOS expression. Notably, P3CSK4 and IFN-β triggered comparable levels of iNOS expression to MRSA infection, while P3CSK4 and IFN-γ led to greater iNOS expression. Next, we infected *Ifnarl-/-* or *Tlr2/4/9-/-* macrophages with MRSA and measured iNOS expression by immunoblot **(Fig 3D, Fig S3F-G).** Deficiency in either signaling pathway abrogated iNOS expression, therefore we conclude that both IFNAR1 and TLR signaling are required for iNOS induction during MRSA infection.

**Figure 3.**
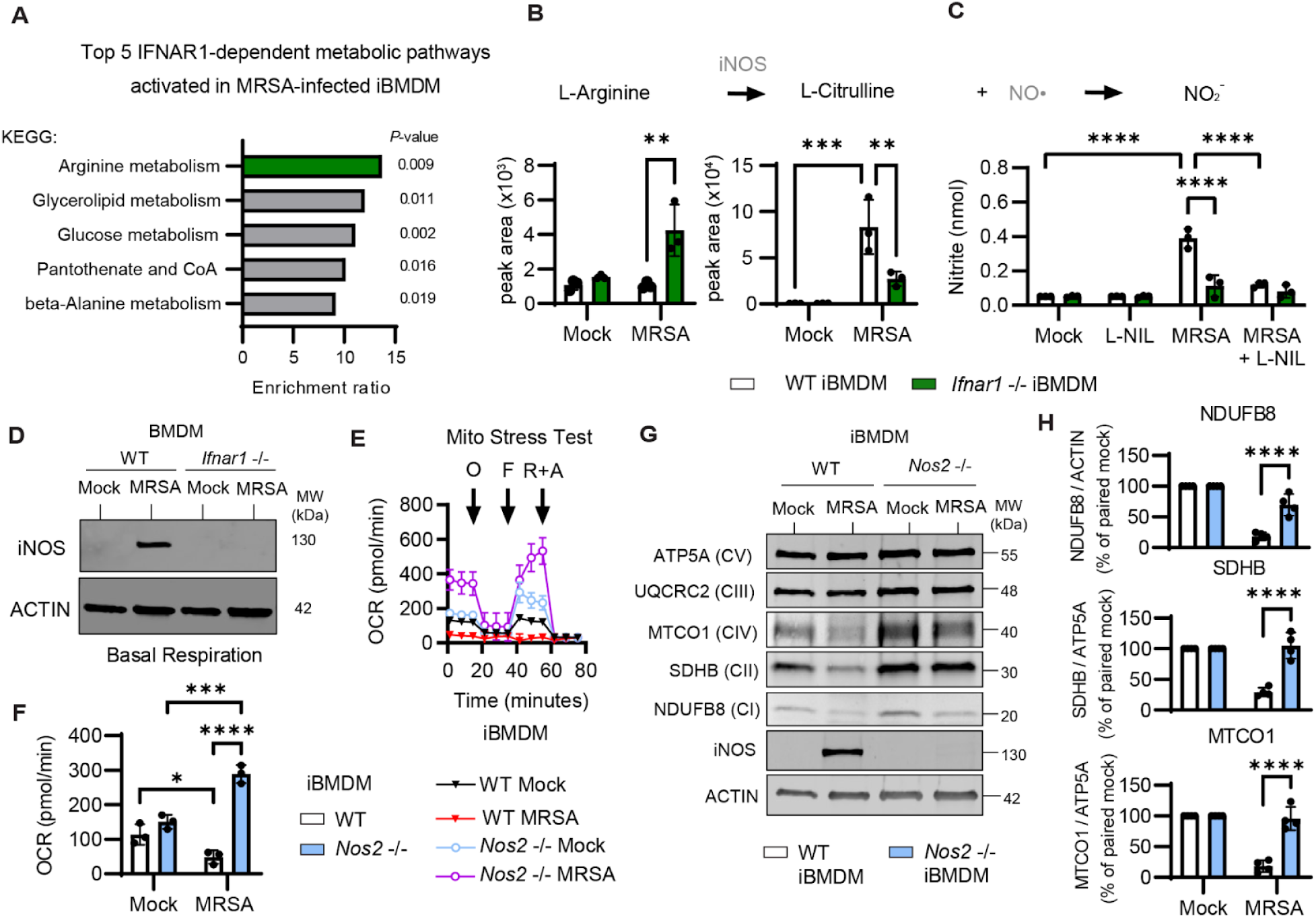
Nitric oxide disrupts the electron transport chain in MRSA-infected macrophages. **A.** LC-MS-based targeted metabolomics analysis of 230 metabolites in mock or 24 h MRSA-infected (MOI 20) WT or *Ifnarl -I-* iBMDM. Metaboanalyst 5.0 pathway enrichment analysis using KEGG annotation of the top 5 significantly changed MRSA-induced metabolic pathways between WT and *Ifnarl -I-* iBMDM. B. Highlighted analysis of peak area from inducible nitric oxide synthase (iNOS)-related metabolites L-arginine and L-citrulline in mock or 24 h MRSA-infected WT and *Ifnarl* -/- iBMDM. C. Griess assay from mock or 24 h MRSA-infected WT or *Ifnarl -I-* iBMDM treated with or without the iNOS inhibitor L-NIL (40 μM). **D.** SDS-PAGE and immunoblot analysis of INOS and ACTIN from in mock or 24 h MRSA-infected WT or *Ifnarl* -/- BMDM whole cell lysates. E. Seahorse XF analysis of the rate of oxygen consumption (OCR) in 24 h MRSA-infected WT and *Nos2* -/- iBMDM using the Mito Stress Test assay, with additions of oligomycin (O), carbonyl cyanide p-trifluoromethoxyphenylhydrazone (FCCP), rotenone (R), and antimycin A (A) at the indicated time points. F. Quantification of basal respiration in mock or 24 h MRSA-infected WT and *Nos2 -I-* iBMDM. **G.** SDS-PAGE and immunoblot analysis of Complex I subunit NDUFB8, Complex II subunit SDHB, Complex III subunit UQCRC2, Complex IV subunit MTCO1, Complex V subunit ATP5A, iNOS, and ACTIN. H. Quantification of the abundance of NDUFB8, SDHB, and MTCO1 relative to ATP5A in mock and MRSA-infected WT and *Nos2 -I-* iBMDM, presented as the percentage of the paired mock sample. Graphs represent the mean of n > 3 biological replicates with SD error bars. P-values were calculated using a two-way ANOVA with Sidak’s post test **(B, C, F,** and **H).** *P < 0.05; **P < 0.01; ***P < 0.001; **‘*P < 0.0001.

Since both iNOS expression and OXPHOS disruption were stimulated by combined TLR and IFNAR signaling, we tested if these processes were functionally related by infecting WT and *Nos2-/-* (iNOS KO) macrophages with MRSA and measuring global metabolic changes with a Seahorse XF assay **(Fig 3E-F).** We found that iNOS was required to dampen respiration during MRSA infection, as well as for loss of Complex I, II and IV RC subunits **(Fig 3G-H).** To validate that NO* had a direct effect on the RCs of the ETC^50^ in our experimental system, we treated BMDM with the NO* donor DETA-NONOate and measured respiration and assembly of the respiratory complexes by Seahorse assay and BN-PAGE, respectively **(Fig S3H-J).** The same RCs lost during MRSA infection were also lost during NO* stimulation. Together, our data demonstrate that TLR and IFN signaling drive iNOS expression, which controls OXPHOS disruption during MRSA infection through NO*-mediated RC disassembly.

### Nitric oxide modulates OXPHOS to shift macrophage cytokine production towards Type I IFN

Having identified the regulatory circuit underlying MRSA-induced OXPHOS remodeling, we then tested how iNOS affects secretion of pro-inflammatory cytokines IL-β, TNF-α, and IFN-β **(Fig 4A).** We found that iNOS deletion in MRSA-infected macrophages enhanced IL-β and TNF-α production, but prevented subsequent IFN-β production. We hypothesized that NO* biased macrophage inflammatory responses towards type I interferon through regulation of OXPHOS and not by microbicidal effects of NO* on MRSA. To test this hypothesis, we pretreated BMDM with the NO* donor DETA-NONOate for 24 h, washed away the NO* donor, infected with MRSA, and then measured secreted levels of IL-β, TNF-α, and IFN-β **(Fig 4B).** NO* suppressed IL-β and TNF-α, but enhanced IFN-β. To determine if RC inhibition phenocopied NO*-dependent effects on cytokine production, we treated BMDM with sub-cytotoxic concentrations of different RC inhibitors, infected with MRSA, and measured secretion of IL-β, IFN-β, and TNF-α **(Fig 4C** and **S4A-B).** Complex I, II, or III inhibition suppressed IL-β, and Complex I or III inhibition enhanced IFN-β. Surprisingly, RC inhibition did not affect TNF-α secretion, indicating that NO* may regulate its expression by another mechanism **(Fig S4B).** Thus, interruption of OXPHOS biases the pro-inflammatory cytokine response to MRSA infection away from IL-ip towards IFN-β **(Fig S4C).**

**Figure 4.**
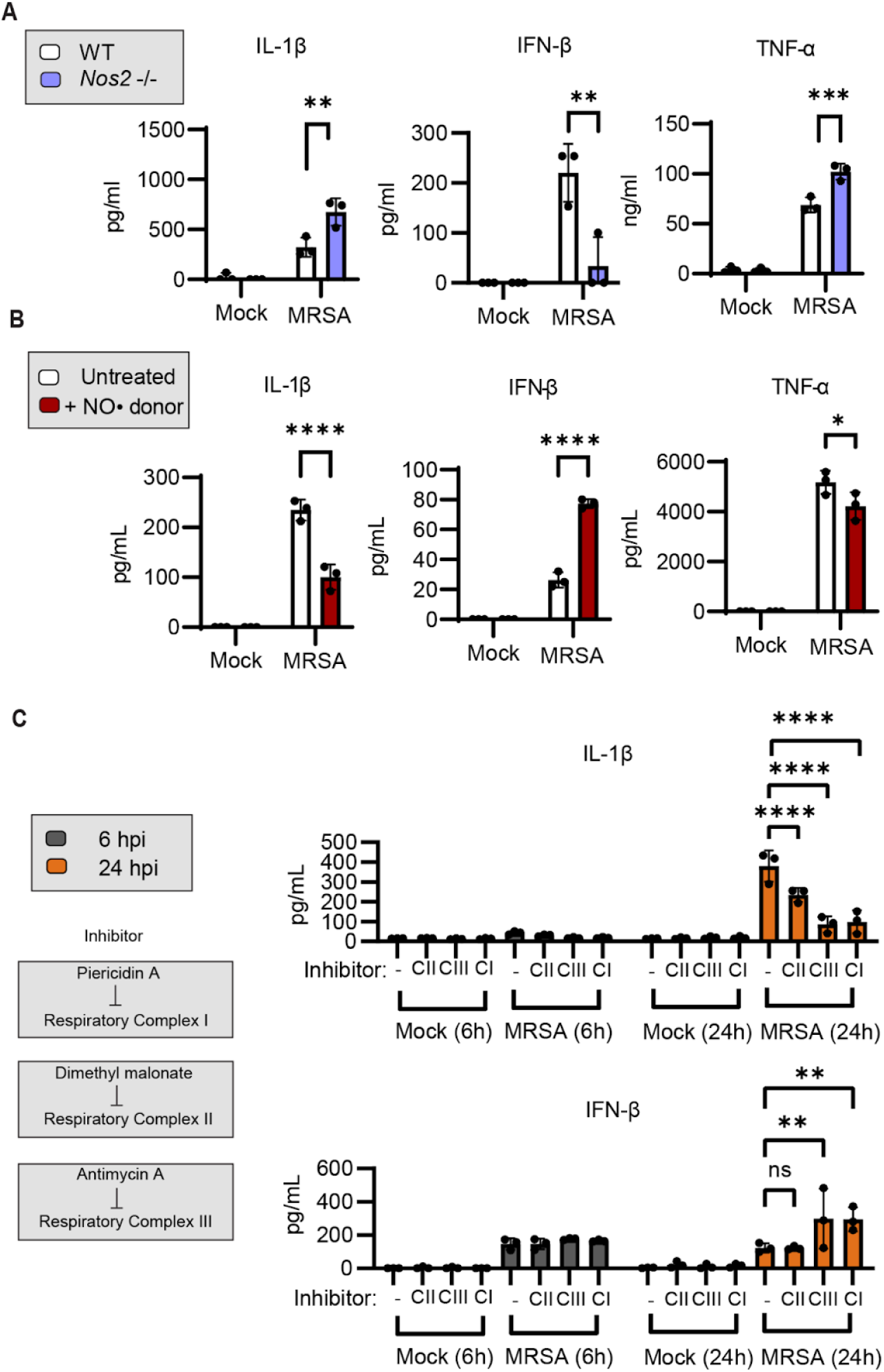
Nitric oxide biases macrophage cytokine production toward type I interferon during MRSA infection. **A.** Enzyme-linked immunosorbent assay (ELISA) analysis of secreted IL-10, IFN-0, and TNF-α from mock or 24 h MRSA-infected WT and *Nos2* -/- BMDM. **B.** ELISA analysis of secreted IL-10, IFN-0, and TNF-α from WT BMDM treated with or without the nitric oxide (NO*) donor DETA-NONOate (500 μM) for 24 h, washed with fresh media, and then mock or MRSA infected for 24 h. **C.** ELISA analysis of secreted IL-10 and IFN-0 after 6h or 24 h MRSA infection with or without sub-cytotoxic doses of respiratory complex inhibitors, dimethyl malonate (Complex II 13 inhibitor; 10 mM), Antimycin A (Complex III inhibitor; 1 μM), and Piericidin A (Complex I inhibitor; 100 nM). Graphs represent the mean of n = 3 biological replicates with SD error bars. P-values were calculated using a two-way ANOVA with Sidak’s post test **(A-C).** *P < 0.05; **P < 0.01; ***P < 0.001; ****P < 0.0001.

### IFN-conditioning limits lactate production and histone lactylation to sustain iNOS expression during MRSA infection

Multiple autoimmune diseases are characterized by chronic expression of type I IFN, which alters the function of immune cells, including macrophages^52^. Furthermore, viral infections, which robustly stimulate type I IFN, are often complicated by bacterial superinfection, suggesting that conditioning of the immune system by type I IFN may deviate productive anti-bacterial responses^40,53,54^. Thus, we reasoned that prior IFN exposure could alter iNOS-mediated metabolic remodeling in MRSA-infected macrophages. We first measured iNOS expression by immunoblot during MRSA infection under naive conditions or after 24 h of conditioning with IFN-β **(Fig 5A-B).** In naive macrophages, iNOS expression peaked at 24 hpi and was sharply suppressed by 48 hpi. In contrast, IFN-β conditioning sustained high levels of iNOS expression at 48h. *Nos2* expression is controlled by epigenetic regulation, and previous studies point to lactate, via histone lactylation, as a mechanism to integrate metabolic changes and transcriptional programming^55,56^. This led us to the hypothesis that chronic type I IFN exposure suppresses lactate production during MRSA infection and prevents lactate-dependent epigenetic repression of iNOS expression. To test our hypothesis, we measured lactate production and histone lactylation after 24 h MRSA infection under naive conditions or after 24h conditioning with IFN-β. IFN-β conditioning inhibited extracellular lactate production in MRSA-infected macrophages **(Fig. 5C).** We also infected naive or IFN-conditioned macrophages with MRSA, or treated cells with lactate directly, and measured H3K18-Lac by IFA and high-content imaging **(Fig 5D-E).** MRSA infection in naive macrophages increased H3K18-Lac levels in the nucleus, which was suppressed by IFN conditioning. In IFN-conditioned macrophages, only lactate supplementation, but not MRSA infection, rescued H3K18-Lac levels. Thus, IFN-conditioning prior to infection decreases macrophage lactate production and histone lactylation. Based on these findings, we predicted that manipulating intracellular lactate levels would alter iNOS expression. Therefore, we treated naive or IFN-conditioned MRSA-infected macrophages with the lactate dehydrogenase (LDH) inhibitor, sodium oxamate, or added lactate directly **(Fig 5F-G).** Inhibition of LDH increased iNOS expression at 48 hpi, while lactate supplementation suppressed iNOS levels, even after IFN-conditioning. We also tested the effect of IFN-conditioning on cytokine production with or without added lactate. IFN-conditioning strongly biased MRSA-infected macrophages towards further IFN-β production and suppressed IL-β **(Fig 5H).** Lactate supplementation reversed the effect of IFN-conditioning up to 48 h post-MRSA infection **(Fig 5I).** Taken together, these data support a model of macrophage infection where type I IFN following TLR signaling upregulates iNOS expression, which shuts down respiration and subsequently enables lactate-dependent epigenetic silencing of *nos2,* providing built-in negative feedback regulation 14 for iNOS-driven metabolic and inflammatory programming. Triggering type I IFN prior to bacterial infection perturbs these immunometabolic checkpoints and dysregulates inflammatory programming.

**Figure 5.**
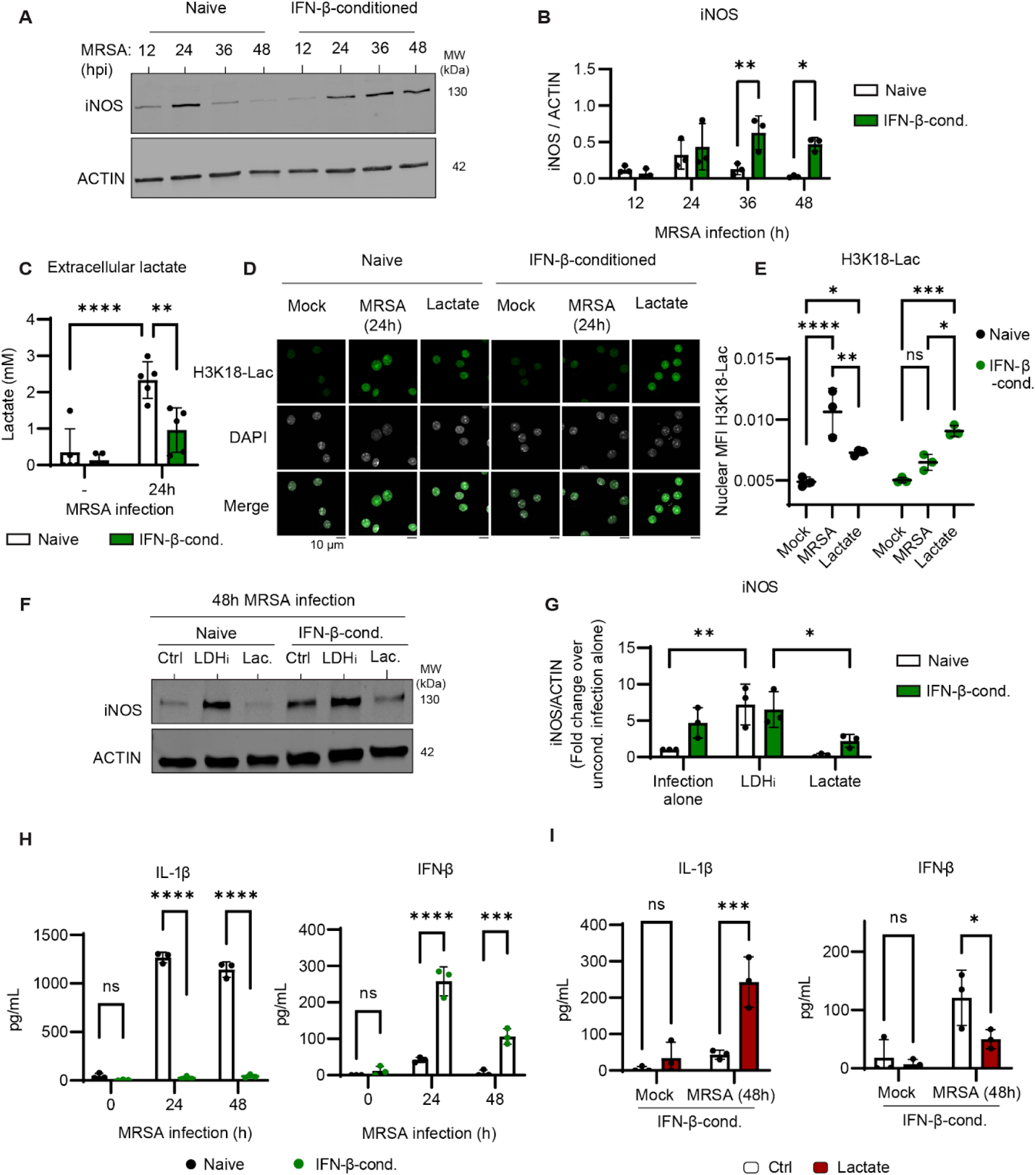
Chronic type I interferon short-circuits macrophage metabolic programming by preventing lactate-mediated iNOS repression. **A.** Immunoblot analysis of iNOS and ACTIN from naive or 24 h IFN-p-conditioned (400U/mL) BMDM after 12, 24, 36, or 48 h of MRSA infection (MOI 20). **B.** Quantification of iNOS relative to ACTIN across multiple experiments. **C.** Measurement of extracellular lactate in naive or 24 h IFN-P-conditioned BMDM after 24 h of MRSA infection. **D.** Automated **15** confocal microscopic analysis of H3K18-Lac IFA from naive or 24 h IFN-p-conditioned BMDM after 24 h of MRSA infection or 24 h lactate (10 mM) treatment. E. Automated quantification of the average nuclear H3K18-Lac intensity per cell using CellProfiler. F. Immunoblot analysis of iNOS and ACTIN from naive or 24 h IFN-p-conditioned BMDM following 48 h MRSA infection in the presence or absence of supplemented lactate (10 mM) or lactate dehydrogenase inhibitor sodium oxamate (LDH;; 10 mM) G. Quantification of iNOS relative to ACTIN represented as the fold change compared to naive macrophages infected with MRSA for 48 h. H. ELISA analysis of secreted IL-β and IFN-β from naive or 24 h IFN-P-conditioned BMDM that were mock or MRSA infected for 24 or 48 h. I. ELISA analysis of secreted IL-β and IFN-β from 24 h IFN-p-conditioned BMDM after 48 h of MRSA infection in the presence or absence of supplemented lactate. Graphs represent the mean of n > 3 biological replicates with SD error bars. P-values were calculated using a two-way ANOVA with Sidak’s post test **(A, C, E, G, H,** and **I).** *P < 0.05; **P < 0.01; ***P < 0.001; ****P < 0.0001.

### Chronic type I interferon in the skin exacerbates iNOS expression, increases inflammation, impairs wound healing, and promotes bacterial dissemination

To test if altering the sequence of type I IFN exposure could dysregulate inflammatory signaling during MRSA infection *in vivo,* we used mice that chronically overexpress the keratinocyte-produced type I IFN, IFN-κ (IFN-κ transgenic [Tg] mice), that display no overt inflammatory disease without specific triggers^57,58^. We infected WT and IFN-κ Tg mice cutaneously with USA300 *lux,* a MRSA strain that constitutively expresses firefly luciferase^59,60^, and monitored bacterial burden longitudinally using an *in vivo* imaging system (MS) **(Fig S5A-B).** Both WT and IFN-κ Tg mice cleared local MRSA infectious burden to a comparable extent by D7, yet the IFN-κ Tg mice displayed an increase in wound area and erythema at D3, macroscopic indications of increased inflammation and defective wound healing **(Fig 6A-D).** Histology confirmed macroscopic features of WT vs. IFN-κ Tg perilesional (PL) skin, revealing elevated inflammation in infected IFN-κ Tg ears **(Fig 6E).** IFN-κ Tg mice exhibited larger PL abscesses with higher neutrophil infiltration and vascular disruption in adjacent tissue **(Fig 6F-G).** According to our *in vitro* data, dysregulation of local inflammation may be caused by perturbations in macrophage metabolism and function via overproduction of iNOS. Tissue immunofluorescence staining of PL iNOS levels revealed an increase in iNOS expression in the MRSA-infected IFN-κ Tg mice compared to WT mice **(Fig 6H-I** and **Fig S5C-D).** In parallel, protein extracts of wound and PL skin showed a significant increase in iNOS levels in MRSA-infected IFN-κ Tg skin compared to infected WT skin by immunoblot **(Fig S5E-F).** Notably, the dominant form of iNOS was truncated (70 kDa) relative to the monomeric size (130 kDa) and may represent tissue-specific regulation of iNOS, possibly via calpain cleavage^61^. Overall, chronic expression of type I IFN in the skin results in local overproduction of iNOS in MRSA wound infections.

**Fig 6.**
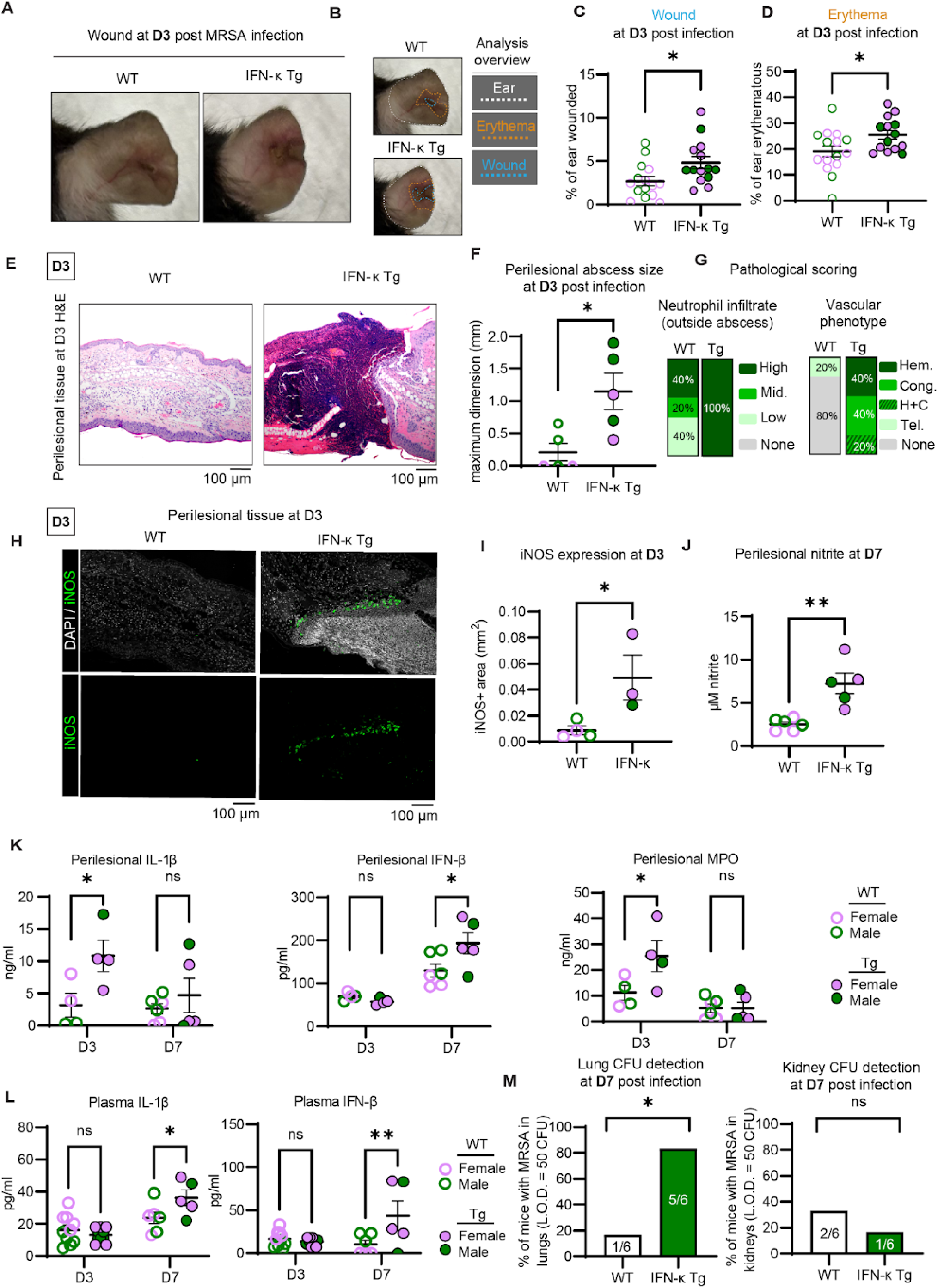
Chronic type I interferon expression in the skin exacerbates iNOS expression, increases inflammation, and impairs wound healing during MRSA infection. Cutaneous MRSA infection in the IFN-κ transgenic (Tg) mouse model which overexpresses IFN-κ in keratinocytes under the *Keratin-14* promoter. Mice were infected with an allergy needle on one ear with MRSA (8e8 CFU; USA300 *lux)* **A.** Photographs of WT and IFN-κ Tg MRSA-infected wound sites. **B.** Analysis overview for assessment of ear, erythema, and wound area. Blinded analysis of wound **(C)** and erythema **(D)** area at day 3 (D3) post-MRSA infection. E. Hematoxylin and eosin (H&E) staining of WT and IFN-κ Tg perilesional (PL) skin sections (sampled from erythematous area) from D3 post-MRSA infection. F. Blinded assessment of PL abscess (neutrophillic inflammation) maximal dimension length as determined by a pathologist. G. Blinded pathological assessment of neutrophil abundance in tissue outside of abscesses, scored as absent, low, intermediate (Mid.), or high and vascular phenotypes where microhemorrhage (Hem.), congestion (Cong.), and telangiectasia (Tel.) were noted. H. Immunohistofluorescence analysis of iNOS protein in PL skin of MRSA-infected WT and IFN-κ Tg mice at D3 post infection. **I.** Quantification of total INOS+ area across PL tissue sections. **J.** Nitrite detection by Griess assay from wound and PL skin extracts of WT and IFN-κ Tg mice at D7 post-infection. K. ELISA analysis of PL and wound extract IL-10, IFN-0, and MPO at D3 and D7 post-infection. L. ELISA analysis of plasma IL-10 and IFN-0 at D3 and D7 post-infection. M. CFU detection above a threshold limit of detection (L.O.D.) value (>50 CFU) from mechanical homogenates from lungs or kidneys at D7 post-infection from WT and IFN-κ Tg mice. Male IFN-κ Tg and WT mice are color-coded in solid green circles and empty green circles respectively, and female IFN-κ Tg and WT mice are color-coded in solid purple and empty purple circles respectively. For wound quantification, graphs represent the mean of n > 15 mice across 2 independent experiments with standard error of the mean (SEM) error bars. Other measurements represent the mean of > 4 mice per genotype with SEM error bars. P-values were calculated using an chi-squared test for binary results **(M),** unpaired t test (C, D, F, I, and J), or two-way ANOVA with Sidak’s post test for grouped analysis (K and L) *P < 0.05; **P < 0.01

After identifying a pronounced iNOS signature in MRSA-infected IFN-κ Tg mice, we measured the key inflammatory signaling molecules IL-β and IFN-β in wound and PL skin across the course of infection. Consistent with our *in vitro* data, IFN-β levels were high at D7 post-infection in IFN-κ Tg wound and PL skin in conjunction with high levels of nitrite **(Fig. 6J-K).** Further, PL skin-associated IL-β was increased in the IFN-κ Tg mice compared to WT at D3 and resolved by D7. A major source of IL-β during MRSA skin and soft tissue infection is neutrophils^62^. Therefore, total recruitment of neutrophils to the infection site was measured using the abundance of the neutrophil-specific enzyme myeloperoxidase (MPO) in wound and PL skin. MPO was increased at D3 in IFN-κ Tg mice compared to WT, indicating increased neutrophil recruitment. By day 7, MPO levels were normalized between WT and IFN-κ Tg PL skin. Thus, our data support unrestrained neutrophil recruitment to the site of infection as a potential source of the early increase in local IL-β. Next, we measured plasma levels of IL-β and IFN-β at D3 and D7 post infection **(Fig 6L).** Systemic IL-β and IFN-β were elevated in IFN-κ Tg mice compared to WT controls at D7 post infection. The observation that systemic cytokines were higher in MRSA-infected IFN-κ Tg mice together with increased vasculopathy, compared to WT mice, led us to hypothesize that bacterial dissemination might occur more frequently in the Tg mice. At D7 pi, we found more frequent bacterial colonization of the lungs, but not the kidneys, of MRSA-infected IFN-κ Tg mice compared to WT mice **(Fig 6M).** Our in vivo data demonstrate that pre-existing constitutive epidermal type I IFN production dysregulates inflammatory responses to MRSA skin infection, preventing wound repair and restriction of bacteria to the site of infection.

## Discussion

In this study, we aimed to identify metabolic drivers that coordinate dynamic macrophage responses to *Staphylococcus aureus* infection. We demonstrate that TLR signaling increases mitochondrial respiration, while the subsequent combined TLR and IFN signaling inputs set off an immunometabolic checkpoint by promoting iNOS expression, leading to OXPHOS disruption and a shift to glycolysis. OXPHOS disruption biased macrophage cytokine production away from IL-β, and towards type I IFN. Accumulation of lactate resulting from accelerated glycolytic flux established a second immunometabolic checkpoint, when it reached a sufficient concentration to repress iNOS expression via histone lactylation. Lactate-mediated iNOS repression then suppressed type I IFN, limiting the kinetics and magnitude of inflammation. Surprisingly, treating macrophages with type I IFN prior to infection, dysregulated this ordered sequence of immunometabolic programming. These results were unexpected because type I IFN is commonly produced by macrophages in response to bacterial infection^26^. Our findings highlight the relevance of evaluating innate immune signaling mechanisms in the context of a natural infection. Importantly, these data establish type I IFN as a pacesetter for these immunometabolic checkpoints during bacterial infection both *in vitro* and *in vivo.* Our results are consistent with a model where IFN-dependent metabolic control of macrophage function coordinates the delicate balance and timing of pro- and anti-inflammatory programming that facilitates host defense and minimizes host damage.

Both bacterial and host-derived molecular pattern recognition and signaling contribute to bacterial clearance, inflammation regulation, and wound healing during MRSA skin infection^38,62–66^. In the early period of infection, when bacterial burden is high, microbial pattern recognition-dominated innate immune signaling can promote a pro-inflammatory environment. In this context, TLR2 heterodimer recognizes bacterial lipoprotein, endosomal TLR9 recognizes bacterial DNA, and the cGAS-STING pathway recognizes cytosolic cyclic dinucleotides, bacterial DNA, or mitochondrial DNA^67,68,69^. This early phase of bacterial pattern recognition results in the secretion of a complex mixture of signaling molecules which are broadly immunomodulatory but calibrated towards robust inflammation. We show that macrophages experiencing an inflammatory environment dominated by TLR signaling adopt a hyper-energetic state characterized by high levels of OXPHOS and glycolysis. Based on prior studies, we speculate that this early hyper-energetic state may support rapid gene expression and produce secondary messengers such as metabolic intermediates or mitochondrial reactive oxygen species (ROS), which can contribute to inflammasome activation^56,70,71,72^. As infection progresses, the high concentration of host-derived signaling molecules changes the inflammatory context of the MRSA-infected lesion^73–75^. In addition to recruiting peripheral leukocytes, including bactericidal neutrophils, host-derived signaling molecules further refine macrophage programming to allow better orchestration of local immunity^26,39^ ^76–78^. Our data are consistent with a recent study indicating an essential role for IFNAR in macrophage metabolic remodeling during *Mycobacterium tuberculosis (Mtb)* infection^15^. We consider the possibility that temporal coordination of inflammation, such as we have described here, is central to responding to the specific context of a given infection, e.g., low or high bacterial burden or low or high levels of host damage due to wounding. Moreover, distinct mechanisms by which specific pathogens engage or modulate the host TLR or IFN signaling pathways may contribute to observed differences in the timing and magnitude of inflammation.

Modulation of mitochondrial metabolism is important for controlling macrophage inflammatory programming^22^. While it is common to assess flux through major metabolic pathways, namely OXPHOS and glycolysis, more detailed metabolite analysis is starting to reveal specific regulatory mechanisms. Notably, recent work highlights a requirement for the ETC in NLRP3 inflammasome activation, IL-β expression and maturation^70,72,79^. In this context, the mitochondrial phospho-creatine pool, derived from OXPHOS, was identified as an important metabolic contributor to NLRP3 inflammasome activation. Since the NLRP3 inflammasome supports MRSA-induced IL-β production, it is possible that depletion of phospho-creatine by OXPHOS shutdown during MRSA infection contributes to IL-β suppression at later stages of infection^80,81^. While phospho-creatine levels were at the limit of detection in our metabolomics samples, we did observe that MRSA infection increased the pool of non-phosphorylated creatine. Future investigation of how OXPHOS disruption during MRSA infection controls cytokine production may reveal important principles underlying the crosstalk between mitochondrial metabolism and innate immunity. Similarly, lactate production is a predictable consequence of the infection-induced glycolytic shift. However, lactate, traditionally considered a waste product of glycolysis and fermentation, accumulates at the site of bacterial infection and now is appreciated as a regulator of antibacterial immunity^56,82^. Indeed, recent evidence suggests that lactate has many roles in the cell, including serving as an alternative fuel source by being converted back into pyruvate by LDH, regulating enzyme activity through post-translational modification, and controlling gene expression epigenetically by modifying histones^56,83–85^. During MRSA infection, host and S. aureus-derived lactate contribute to lactate buildup at the sites of infection. Our data support that lactate accumulation at this later stage in infection enables iNOS repression through histone lactylation. Of note, the *Nos2* promoter exhibits high levels of histone lactylation during inflammatory macrophage activation^56^, and lactate was recently shown to decrease *Nos2* expression in *Salmonella* Typhimurium-infected macrophages^86^. Consistent with its inhibitory effects on iNOS expression, we report that lactate suppresses type I IFN production in MRSA-infected macrophages. Thus, we propose that late-stage infection-associated lactate accumulation acts as a metabolic checkpoint that prevents NO*-mediated chronic type I IFN-biased cytokine responses.

Conditions that elicit robust type I IFN production, such as certain autoimmune diseases, inflammatory diseases, or viral infections, tend to exacerbate the severity of MRSA infection^30,40^. Consistent with our in vitro IFN-conditioning experiments, mice that consitutively express epidermal IFN-κ (IFN-κ Tg) prior to cutaneous MRSA infection exhibited impaired wound healing and poor resolution of local inflammation compared to WT controls. These results suggest short-circuiting of the metabolic sequence orchestrated by type I IFN during MRSA infection may contribute to the defect in wound healing we observe in the IFN-κ Tg mice. This result is in contrast to diabetic mice where IFN-κ aids in wound healing^58^. Similarly, although iNOS contributes to healing in some wound models^87^, overproduction of iNOS in the highly inflammatory environment of a MRSA-infected wound may impair key aspects of macrophage metabolic regulation required for effective healing^87,88^. This notion is supported by the strict kinetic regulation of iNOS and its function in MRSA-infected macrophages in determining pro-inflammatory cytokine output, especially IFN-β. Moreover, in addition to exacerbated skin inflammation, in our infection model, IFN-κ Tg mice exhibited increased bacterial infection in the lung. The vascular damage observed around the MRSA-infected wound in IFN-κ Tg compared to WT mice suggests a possible hematogenous route of bacterial spread to the lungs. Notably, type I IFN has been shown to enhance vascular inflammation in autoimmune disease^52,89,90^.

Overall, our findings suggest a requirement for ordered timing of inflammatory and metabolic signals that guide the progression of inflammation during MRSA infection. These results point to type I IFN as a critical regulator that coordinates iNOS-dependent OXPHOS shutdown following a TLR-driven increase in respiration during MRSA infection. Conditions that sustained iNOS expression further enhanced type I IFN production in a positive feedback loop reminiscent of chronic diseases of IFN overexpression, such as systemic lupus erythematosus^91^. Lastly, we identify lactate-mediated iNOS repression as a metabolic checkpoint that is impaired in conditions that mimic chronic interferon signaling. Overall, our findings define type I IFN as a key temporal determinant of immunometabolic programming during MRSA infection.

## Methods

### Animal use

Experimental animals were housed at the University of Michigan Medical School Unit for Laboratory Animal Medicine (ULAM) in climate-controlled, specific pathogen-free facilities and treated humanely in accordance with an Institutional Animal Care Use for Research Committee-approved protocol (PRO00010463).

### Cell culture

Murine primary bone marrow-derived macrophages (BMDM) and immortalized BMDM (iBMDM) were prepared as previously reported. Femurs and tibiae from the C57BL/6J lineage mice of various genotypes were flushed and grown for 6 days in macrophage differentiation media [50% DMEM, 1 mM sodium pyruvate, 2 mM L-glutamine, 20% heat-inactivated fetal bovine serum (FBS) (Biowest), 30% L929 cell-conditioned medium, penicillin (50 U/ml), and streptomycin (50 pg/ml)]. For immortalization, recombinant Cre-J2 virus containing v-Raf and v-Myc oncogenes was generated in 3T3 fibroblasts grown in Dulbecco’s modified Eagle’s medium (DMEM) supplemented with 10% heat-inactivated FBS and penicillin (50 U/ml) and streptomycin (50 pg/ml). Sterile-filtered culture supernatants containing Cre-J2 virus were stored at -80°C. Bone marrow isolates were transduced with Cre-J2 virus in macrophage differentiation media to generate iBMDMs, and iBMDMs were grown for at least 1 month to ensure successful immortalization. L-929 cells were cultured in minimum essential Eagle’s medium supplemented with 1 mM nonessential amino acid, 1 mM sodium pyruvate, 2 mM l-glutamine, 10 mM Hepes, and 10% FBS. L929-conditioned medium was sterile filtered prior to use. All experiments were performed in DMEM supplemented with 2 mM l-glutamine, 1 mM sodium pyruvate, and 10% FBS, without antibiotics unless otherwise indicated. *Nos2* -/- (strain #002609), *Ifnarl* -/- (strain #010830), and wild-type (WT) age and sex-matched pairs were obtained from the Jackson Laboratory. BMDMs were derived according to the protocol described above. Effective iNOS knockout was validated by immunoblot with a murine iNOS-specific antibody (1:1000; Cell Signaling Technology, 2982S). *Tlr2/4/9 -I-* iBMDM were previously generated in our laboratory from mice provided by Dr. Tod Merkel and have been validated using isoform alignment from RNA sequencing data and cytokine production **(Fig S6A-B).** *Ifnarl -I-* deletion was validated using PCR with primer sequences provided by the Jackson laboratory 22 (5’-CGA GGC GAA GTG GTT AAA AG-3’, 5’-ACG GAT CAA CCT CAT TCC AC-3’, and 5’-AAT TCG CCA ATG ACA AGA CG-3’) **(Fig S6C).** Cell lines were confirmed to be mycoplasma-free with the Lookout Mycoplasma PCR Detection Kit (Sigma-Aldrich). Genotyping of various mice used in this work was accomplished with the APExBIO direct mouse genotyping kit and primers described in **Table S1.** When indicated, macrophages were treated with the TLR2 agonist PamCSK4, abbreviated P3CSK4 (2 pg/mL; Invivogen), or recombinant IFN-β (400 U/mL; Pestka Biomedical Laboratories) or IFN-γ (50 ng/mL; Peprotech). Chemicals and inhibitors were used in cell culture, and the origin and concentration of each chemical are appended in **Table S2.** All cells were cultured at 37°C in 5% CO2.

### MRSA infection of macrophage cultures *in vitro*

Methicillin-resistant *Staphylococcus aureus* (MRSA) USA300 isolate JE2 glycerol stocks were struck out on tryptic soy broth (TSB) agar plates within 3 days of planned experiments. For each experiment, a single MRSA colony was picked and grown overnight at 37°C, slanted, with shaking. Overnight cultures were washed three times with PBS and diluted to an optical density (OD) of 1 as measured by a spectrophotometer, which was determined to equate to a bacterial density of 10^9^ CFU/mL. Macrophages were infected at a multiplicity of infection (MOI) of 20 CFU/macrophage and incubated at 37°C in a tissue culture incubator for 1 h. After 1 h, cultures were supplemented with 10 U/mL of lysostaphin (Sigma) to kill extracellular bacteria. Infections were then allowed to proceed to the endpoint according to individual experimental protocols.

### Seahorse extracellular flux (XF) assay

The rate of oxygen consumption (OCR) and the rate of extracellular acidification (ECAR) of macrophages in culture was measured using an Agilent Seahorse XF96 analyzer. Macrophages were plated in a Seahorse 96-well assay plate and cultured overnight in DMEM supplemented with 10 mM glucose, 1 mM pyruvate, 2 mM glutamine, and 10% FBS (stimulation medium). A target cell density of 80% confluence was achieved by addition of 35,000 iBMDM or 45,000 BMDM per well. On the next day, the medium was replaced with fresh stimulation medium with or without MRSA (MOI 20), P3CSK4 (2 pg/mL), IFN-β (400 U/mL), or DETA-NONOate for 24 h. After 6 hours, medium was exchanged for DMEM supplemented with the same levels of glucose, pyruvate, and glutamine without FBS. Cells were kept in a 37°C incubator without CO2 for 30 min before analysis. In all experiments, the Mito Stress Test kit from Agilent was used to probe different aspects of mitochondrial function with respiratory chain inhibitors, 1.5 μM oligomycin, 2 μM carbonyl cyanide p-trifluoromethoxyphenylhydrazone (FCCP), 0.5 μM rotenone, and 0.5 μM antimycin A. FCCP was titrated in a pilot experiment to determine optimal induction of maximal respiration. Comparable plating between conditions was confirmed by measuring protein content by Bradford assay (Bio-Rad) from a replicate 96-well plate. As we have observed that cell density does not correlate linearly with extracellular flux measurements, we did not normalize OCR or ECAR measurements. Instead, experiments were maintained with cellular confluence within a threshold of 20% between experimental conditions.

### Targeted metabolomics

WT BMDM derived from five different C57BL/6J mice, 2 male and 3 females, (1e6 cells) were infected with MRSA (MOI 20) for 24 h or left uninfected. Alternatively, WT and *Ifnarl* -/- iBMDM derived from three different mice per genotype, 2 males and 1 female, were infected with MRSA (MOI 20) for 24 h or left uninfected. All experiments were performed in DMEM supplemented with 2 mM l-glutamine, 1 mM sodium pyruvate, and 10% FBS. At the experimental endpoint, cells were washed with ice cold PBS and metabolites were extracted with 80% methanol for 10 min on dry ice. Metabolite extracts were centrifuged at 16,000 g for 10 min at 4°C to remove insoluble material, and the supernatant was collected. Extracts were protein normalized by adjusting volumes according to a Bradford assay from paired samples for each experiment. Equivalent volumes of normalized extracts were dried using a SpeedVac at room temperature (RT) for 2.5h. After drying, the metabolite pellet was dissolved in 50% methanol-water and analyzed by liquid chromatography-mass spectrometry (LC-MS) with an Agilent 1290 Infinity II LC-6470 Triple Quadrupole (QqQ) tandem mass spectrometer system in the negative mode. Agilent ZORBAX RRHD Extend’d8, 2.1 x 150 mm, 1.8 μM and ZORBAX Extend Fast Guards for UHPLC were used in separation. The LC gradient utilized 3 solutions: Solution Awas 97% water and 3% methanol with 15 mM acetic acid and 10 mM Tributylamine (TBA) at pH 5. Solution B was 15 mM acetic acid and 10 mM TBA in methanol. Washing Solvent C was acetonitrile. The following LC gradient profile was used: from 0-2.5 min, 0.25 ml/min of 100% A; from 2.5-7.5 min, 0.25 ml/min of 80% A and 20% B; from 7.5-13 min, 0.25 ml/min of 55% A and 45% B; from 13-20 min, 0.25 ml/min of 1% A and 99% B; from 20-24 min, 0.25 ml/min of 1% A and 99% B; from 24.05-27 min, 0.25 ml/min of 1% A and 99% C; from 27.5-31.35 min, 0.8 ml/min of 1% A and 99% C; from 31.5-32.25, 0.6 ml/min of 1% A and 99% C; from 32.25-39.9 min, 0.4 ml/min of 1% A and 99% C; from 39.9-40 min, 0.25 ml/min of 100% A. Targeted metabolomics analysis of measured MS/MS spectra with a reference library of 230 standard metabolites was performed using Skyline open-source software, and major findings were validated by an independent assessment using the Agilent Masshunter Workstation Software LC/MS Data Acquisition for 6400 Series Triple Quadrupole MS with Version B.08.02. The LC-MS/MS dynamic multiple reaction monitoring (dMRM) method for detection of 220 metabolites based on MS/MS spectra, retention time (RT) windows, and explicit RTs was developed by Agilent using 220 individual metabolite standards^51^’^92^ ^93^. An additional 10 metabolites were added to the acquisition method using an identical chromatographic method with compound standards. 0.07-min peak width was used in dMRM scans with an acquisition time of 24 min. Significantly changed metabolites between mock and MRSA-infected WT BMDM or MRSA-infected WT iBMDM and MRSA-infected *Ifnarl -I-* iBMDM were determined using unpaired t-tests and filtering of *P* < 0.05 and absolute value of log_2_(fold change) > 0.5. Metabolites with low abundance (<1000 peak area) or poor signal to noise ratio (<3) compared to matrix blank were omitted from analysis. Pathway enrichment analysis (Kyoto Encyclopedia of Genes and Genomes; KEGG) was performed using Metaboanalyst 5.0, filtering the top 5 pathways (P-value) from each group and sorting based on enrichment score.

### Flow cytometric analysis of mitochondrial membrane potential and cell viability

WT BMDM or WT and *Hr2/4/9* -/- iBMDM were infected with MRSA (MOI 20), left unstimulated, or treated with P3CSK4 (2 pg/mL) with or without IFN-β (400 U/mL) or IFN-γ (50 ng/mL). After 24 h stimulation, cells were stained with a cocktail of MitoTracker Deep Red (MTDR; 50 ng/mL) and MitoTracker Green (MTG; 50 ng/mL) or propidium iodide (PI; 2 pg/mL) alone for 30 min at 37°C. Cells were washed twice in fluorescence-activated cell sorting (FACS) buffer (PBS + 10% FBS) and analyzed with a flow cytometer (Fortessa, BD Biosciences) with unstained and fluorescence minus one control (FMO). Analysis was performed using FlowJo software.

### Denaturing protein extraction, SDS-PAGE, and immunoblot

Cells were washed with ice-cold PBS and then lysed in 1% NP-40 lysis buffer (50mM Tris-HCI [pH 7.4], 150 mM NaCI, 1% IGEPAL CA-630 [Sigma]) supplemented with Halt Protease and phosphatase inhibitors for 15 min on ice (Thermo Fisher Scientific). Prior to sodium dodecyl sulfate-polyacrylamide gel electrophoresis (SDS-PAGE), sample protein content was normalized by dilution following a Bradford assay. Samples were diluted in Laemmli sample loading buffer supplemented with |3-mercaptoethanol (Bio-Rad), heated for 5 min at 95°C, and then separated on 4 to 20% gradient polyacrylamide tris-glycine gels (Bio-Rad). For respiratory complex subunit immunoblots (IB), samples were instead heated for 20 min at 50°C to prevent precipitation. In the case of H3K18-Lac IB, histones were first acid-extracted using an Abeam histone extraction kit (ab113476). After SDS-PAGE, protein was transferred to a 0.45-μM nitrocellulose membrane (Cytiva) with a Transblot Turbo semi-dry transfer system (Bio-Rad). Membranes were blocked with 5% BSA and 0.1% Tween 20 (IB blocking buffer) for 30 min at RT and then incubated with primary antibody in IB blocking buffer overnight at 4°C. Blots were stained with LI-COR IRdye secondary antibodies and imaged with an Odyssey IR Imager. Occasionally, total protein was visualized by staining gels with Revert700 prior to blocking and imaging with an Odyssey IR Imager (LI-COR). IBs were quantified using the Imaged densitometric gel analysis protocol for 1D gels. Detailed information about antibodies used in this study can be found in **Table S2.**

### Native protein extraction, BN-PAGE, and immunoblot

Native respiratory complex analysis was achieved through blue native polyacrylamide gel electrophoresis (BN-PAGE) and immunoblot, according to established protocols and using commercially available reagents (Thermo Fisher Scientific). Whole-cell extracts (1e6 cells) were solubilized with 2 mg of digitonin in native PAGE sample buffer (Thermo Fisher Scientific, BN2008) for 30 min on ice. Extracts were centrifuged for 10 min at 17,000 g and 4°C to remove insoluble material. 0.5% Coomassie G-250 was added to each soluble fraction immediately prior to loading samples on native PAGE 4 to 16% bis-tris mini gels (Thermo Fisher Scientific). Electrophoresis was performed at 4°C for 30 min in dark blue cathode at 150 V (Thermo Fisher Scientific, BN2007). After 30 min, the cathode buffer was switched to the light blue cathode buffer. Gels were washed with ultrapure water and then soaked with 2* NuPAGE transfer buffer with 0.04% SDS for 15 min (Thermo Fisher Scientific, NP0006). Protein was transferred to an Immobilon-FL polyvinylidene difluoride membrane with 2x NuPAGE transfer buffer supplemented with 10% methanol at 15 V for 15 min using a Transblot Turbo semi-dry transfer system (Bio-Rad). After transfer, membranes were fixed with 8% acetic acid for 5 min. Membranes were washed with water, 100% methanol, and then once again with water to remove excess Coomassie G-250. Immunoblot and downstream analysis were performed as described above.

### IFA and automated confocal microscopy

Macrophages were plated in 96-well or 384-well imaging-grade plates (Phenoplate, Perkin Elmer). At each experimental endpoint, macrophages were fixed with freshly prepared 4% paraformaldehyde (PFA) at RT for 15 min. Immunofluorescence assays (IFA) and imaging were always performed on the day of each experimental endpoint. Wells were washed with PBS + 0.1% Triton X-100 (wash buffer). Wells were blocked with 5% BSA and 10% goat serum in wash buffer (block buffer) for 30 min at RT. A cocktail of primary antibodies was prepared in block buffer and incubated in each well for 1 hour at RT. Wells were washed with wash buffer and incubated with secondary antibodies and counterstains in block buffer at RT for 30 min. Wells were washed with PBS and immediately imaged at40X magnification on a CQ1 or Cell Voyager 8000 automated confocal microscope (Yokogawa). Maximum intensity projection (MIP) images were collected across 8 Z-planes spanning 10 μM and centered around a laser autofocus-defined focal plane for each analysis image. Using this automated method >1000 cells were sampled per condition for each experiment. Specific antibody details are available in **Table S2.**

### Automated image analysis

Open-source image analysis software CellProfiler was used for quantification of all micrographs CellProfiler pipelines related to this manuscript are available in the Supplementary Materials. Image analysis was performed on raw images (MIP images from Yokogawa instrument software) using the University of Michigan Advanced Research Computing Great Lakes computing cluster. Automated single-cell analysis was achieved by segmentation of nuclear objects based on nuclear staining, Hoechst or DAPI (4’,6-diamidino-2-phenylindole), using the identify primary objects module, followed by propagation of the nuclear objects to the cellular periphery based on a whole-cell stain or a empirically defined number of pixels roughly equal to the mean nuclear diameter using the identify secondary objects module. Following cellular segmentation, the intensity of a variety of stains in the total cellular, nuclear, and cytoplasmic area was measured on a single-cell basis. In the case of phospho-STAT1 IFA, the total cellular area was measured. In the case of H3K18-Lac IFA, the nuclear intensity was measured. In the case of NF-κB p65 IFA, the ratio of nuclear to cytoplasmic signal was calculated as a measure of NF-κB nuclear translocation. Removal of outliers at the single cell level was performed using strict ROUT outlier identification (Q = 0.1%) in GraphPad Prism from cell-level data pooled across multiple experiments. For quantification of immunohistofluorescence, the area of the entire tissue section based on the DAPI stain and the iNOS+ area based on the iNOS stain were measured using the identify primary objects module in CellProfiler. Representative images shown in this manuscript have uniformly scaled brightness and contrast within each experiment.

### ELISA

Macrophages were stimulated under various experimental conditions detailed above. At the experimental endpoint, cell culture supernatants were collected and centrifuged to remove cellular debris. Following *in vivo* MRSA infection, peripheral blood was collected by cardiac puncture and plasma was separated using EDTA-coated plasma collection tubes (BD Bioscience). Additionally, PL and wounded skin of MRSA-infected mice was submerged in 1% NP-40 lysis buffer in pH 7.4 TBS supplemented with Halt protease and phosphatase inhibitors (Thermo Fisher Scientific) and ground with a mortar and pestle on ice to generate a wound homogenate. Culture supernatants, plasma samples, and wound homogenates were immediately frozen at -80°C and analyzed within two weeks of sample generation. TNF-α (DY 410, R&D Systems), IL-6 (DY 406, R&D Systems), IL-β (DY 401, R&D Systems), and IFN-β (DY 8234-05, R&D Systems) were measured by enzyme-linked immunosorbent assay (ELISA) by the University of Michigan Cancer Center Immunology Core. Myeloperoxidase (MPO) in tissue was measured using a commercially available kit from Abeam according to manufacturer recommendations (ab155458). To minimize batch effects between experimental replicates, reported supernatant cytokine measurements are mean-centered between experiments.

### Measurement of lactate and nitrite in culture supernatants and tissue

The abundance of lactate (Cayman, 600450) and nitrite (Abeam, ab234044) in culture supernatants were measured using commercially available colorimetric assay kits for these compounds. These assays were performed according to manufacturer recommendations from tissue or cell culture supernatant following macrophage infection with MRSA in the context of WT vs. *Ifnarl -I-* or naive vs. IFN-β conditioning. Tissue extracts were prepared by submerging murine ear skin in 1% NP-40 in pH 7.4 TBS and grinding with an ice cold mortar and pestle. In the case of nitrite, supernatants and tissue extracts were immediately analyzed. For lactate measurements, supernatants were frozen and processed within one week of sample generation.

### Cutaneous MRSA skin infection model

Female and male WT C57BL/6J mice (Jackson Laboratory) and IFN-κ transgenic (Tg) mice, which harbor *Ifnk* under the keratin 14 promoter to sustain *Ifnk* overexpression in the epidermis **(Fig S6D),** aged 8 to 16 weeks, were housed in specific pathogen-free and climate-controlled facilities at the University of Michigan Medical School Unit for Laboratory Animal Medicine. Chow (5L0D) and water were provided to mice *ad libitum.* IFN-κ Tg mice were validated using the following primer sequences *(Ifnk* forward: 5’-AGCTCAAGAGTGCTTCATGGAC-3’, *Ifnk* reverse: 5’-GTATTTGTGAGCCAGGGCATTG-3’ and internal control forward: 5’-CTATCAGGGATACTCCTCTTTGCC-3’, internal control reverse: 5’-GATAACAGGAATGACAAGCTCATGGT-3’). USA300 *lux,* a strain of MRSA with constitutive expression of firefly luciferase, was provided by the laboratory of Dr. Alexander Horswill^94^. USA300 *lux* was struck out from a glycerol stock onto TSA plates two days before liquid culturing. To generate USA300 *lux* liquid cultures, a single colony was picked and grown overnight in TSB at 37°C, slanted, with shaking. On the next day, the overnight culture was split 1:100 in fresh TSB to generate cultures at an OD of 0.8 as measured by spectrophotometer. OD 0.8 cultures in TSB were aliquoted and frozen at -80°C. Frozen OD 0.8 cultures were thawed, diluted 1:4 in fresh TSB, and expanded again to OD 0.8 two hours prior to infection on day 0 (DO) of infection. The expanded OD 0.8 culture was collected by centrifugation, washed three times with PBS and diluted to 1e11 CFU/mL. *In vivo* MRSA infection followed a cutaneous ear infection model which has been previously described^60^. To begin, mice were anesthetized with 3% isoflurane and the right ear was cleaned with 70% isopropanol. Sterile Morrow Brown allergy needles were coated with 8 pL of 10^11^ CFU/mL MRSA *lux* in PBS. Cutaneous injury and MRSA infection on the cleaned ear was achieved by 10 proximal punches with the MRSA-coated allergy needle. Mice were monitored daily with assessment of behavior, weight, bioluminescence signal in the ear, and the macroscopic appearance of the ear. Mice were euthanized at D3 or D7 post-infection as the experimental endpoints.

### Longitudinal bacterial burden measurement

Bacterial burden in MRSA /ux-infected mouse ears was monitored every 24 h for 3 or 7 consecutive days using an In Vivo Imaging System (IVIS) Spectrum (Perkin Elmer). Bacterial burden was quantified as the bioluminescence (radiance) within a fixed region of interest (ROI) centered around the infected ear. Bacterial burden is reported longitudinally as the percentage of bioluminescence signal relative to D1.

### Assessment of ear wound and erythema size

Color images of MRSA /ux-infected ears were taken at D3 post-infection (iPhone 14, Apple). Images were blinded and areas of wounded and red (erythematous) skin were quantified independently by two scientists who were not directly involved in the study. Quantification was performed by outlining the total ear area, wound area, and erythematous area in Imaged. Measurements were averaged across the blinded analyses. Wound and erythematous area are reported as a percentage of total ear area. Removal of outliers was performed using ROUT outlier identification (Q = 1%) in GraphPad Prism which resulted in the exclusion of 1 mouse from each genotype.

### Assessment of bacteria in organs

Mice were euthanized at D3 and D7 post cutaneous MRSA infection and then lungs and kidneys were mechanically homogenized using an automatic bead disrupter. Undiluted homogenized organs were spot plated in triplicate (20 uL; 4% total organ content) for CFU on TSA plates. Bacterial presence was defined as at least 50 CFU in total organ homogenate as previously reported for a similar model, with the limit of detection defined as a mean of 2 CFU across triplicate spots^60^. Bacteria were not detected at D3 post-infection in lung or kidney, so D7 data are presented alone. One IFN-κ Tg male mouse succumbed to infection at D6 post-infection. Necropsy of this mouse revealed bioluminescence signal in the lung but not the kidneys or liver, so this was considered a lung bacterial spread event.

### H&E staining and histopathology scoring

Perilesional (PL) murine skin was fixed in 10% formalin, dehydrated, embedded in paraffin, sectioned, and stained with hematoxylin (Surgipath, 3801540, Leica Biosystems) and eosin (Surgipath, 3801600, Leica Biosystems). H&E staining was performed per standard protocols. Scoring of inflammation was conducted in a blinded fashion by a dermatopathologist (PH). The maximal dimension (length) of PL abscesses was measured. Pathological assessment of neutrophilic inflammation and vascular damage in PL tissue was performed.

### Tissue immunofluorescence

Formalin-fixed, paraffin-embedded specimens of murine PL skin on slides were heated for 60 min at 60 °C, rehydrated, and epitope-retrieved with tris-EDTA (pH 6). Slides were blocked with normal goat serum 5% in PBS, incubated with primary antibody overnight at 4 °C, washed, incubated with secondary antibodies, and counterstained with DAPI. Entire sections were imaged at 20X magnification on a CQ1 automated confocal microscope (Yokogawa). A maximal intensity projection (MIP) of 8 Z-planes spanning 10 μM and centered around a laser-defined focal plane was recorded for each analysis image. Analysis was performed in CellProfiler, by guantifying iNOS+ area (immunofluorescence thresholding). Specifics of antibodies are outlined in **Table S2.**

### RNA-seguencing and analysis

WT and *Tlr2/4/9* -/- iBMDM were lysed in TRIzol reagent (Thermo Fisher) and RNA extraction was performed according to manufacturer protocol. Samples were then treated with Turbo DNase I (Thermo Fisher) and incubated at 37 °C for 30 min, followed by phenol/chloroform extraction and ethanol precipitation overnight. RNA concentrations were then normalized. PolyA enrichment was performed and sequencing libraries were prepared and sequenced on an Illumina NovaSeq 6000 with 150 bp paired-end reads by Novogene. RNA-seq analysis was performed in Galaxy (usegalaxy.org). Reads were evaluated using FastQC and trimmed using cutadapt^95^, followed by quantification of transcripts from the GRCm38 mouse genome using STAR^96^. BAM and BAI files were generated using Galaxy and visualized using the Interactive Genomics Viewer^97^ desktop application.

## Resource availability

### Data and code availability

Upon final publication, raw targeted metabolomics data will be made available through open access on the University of Michigan Deep Blue repository. Processed targeted metabolomics data are available in **File S1** and **File S2.** Cell Profiler pipeline files used for image analysis will be available as supplementary files. All other relevant data are included in this manuscript.

### Materials availability

All unique/stable reagents generated in this study are available from the Lead Contact (M.X.O) without restriction.

## Acknowledgments

We thank Dr. Alexander Horswill for providing the USA300 *lux Staphylococcus aureus* strain and Dr. Tod Merkel for providing *Tlr2/4/9 -I-* mice. We thank Dr. Basel Abuaita and Dr. Marie-Eve Charbonneau for the generation of *Tlr2/4/9 -I-* and *Nos2 -I-* iBMDM. We thank Joel Whitfield in the Rogel Cancer Center Immunology core for routine ELISA analysis. We thank Deborah Colesa for organizing and genotyping the IFN-κ Tg mouse colony. The Rogel Cancer Center Tissue and Molecular Pathology (TMP) Core prepared histological samples. The Advanced Research Computing Great Lakes computing cluster was used for high content image analysis. BioRender software with an academic license was used to generate illustrations for this manuscript.

## Author contributions

M.B.R. and M.X.O. contributed to the conceptualization of the research study. Data curation and formal analysis were completed by M.B.R., B.K., and M.J.M. Funding was acquired by M.X.O., J.M.K., C.A.L., T.R.O., and J.Z.S. Investigation was completed by M.B.R., B.K., N.K.J., H.E.N., B.C.M., T.L.S., D.A., L.Z., and P.W.H. Critical resources and supervision were provided by J.M.K., J.Z.S., P.W.H., and C.A.L. Writing and visualization of the original manuscript draft were completed by M.B.R., B.K., and M.X.O. Review and editing were completed by all authors.

