## Supplemental Figures for "Type I interferon governs immunometabolic checkpoints that coordinate inflammation during *Staphylococcal* infection"

### Figure S1

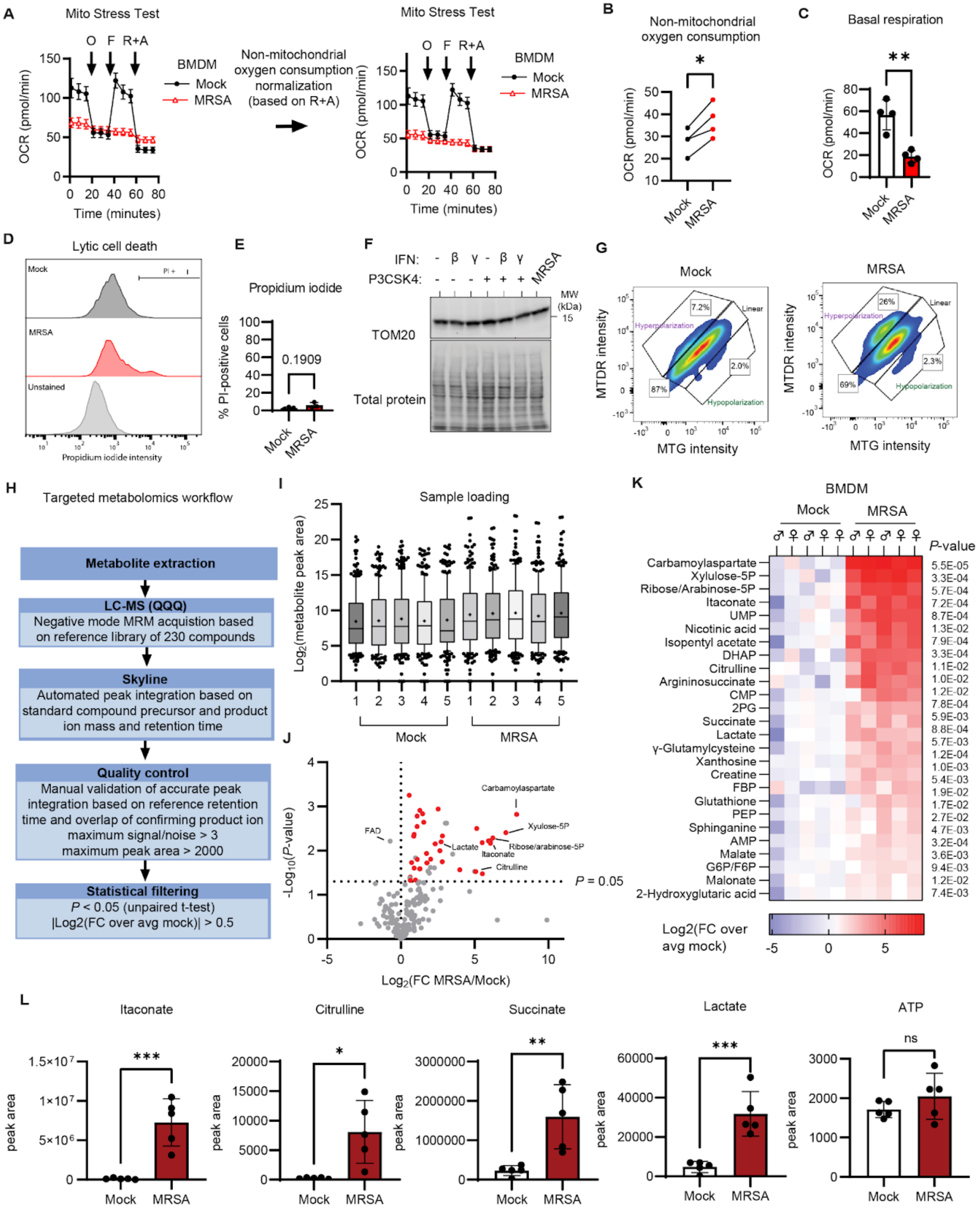

**Fig S1**. **Supplemental data supporting Figure 1. A.** Seahorse XF analysis of the rate of oxygen consumption (OCR) in 24 h MRSA-infected BMDM using the Mito Stress Test assay, with additions of 1.5 μM oligomycin (O), 2 μM carbonyl cyanide p-trifluoromethoxyphenylhydrazone (FCCP), 0.5 μM rotenone (R), and 0.5 μM antimycin A (A) at the indicated time points. Non-mitochondrial respiration (post R+A addition) was normalized between conditions to account for high non-mitochondrial respiration in infected conditions (**B**), and basal respiration (**C**) was calculated from raw data using Agilent Wave software. **D.** Flow cytometric analysis of lytic cell death using propidium iodide (PI) staining at 24 h post infection. **E.** Percentage of PI+ cells across multiple experiments indicates that less than 10% of BMDM in the MRSA-infected condition undergo lytic cell death. **F.** Whole cell extracts from BMDM treated with TLR2 agonist P3CSK4 with or without IFN-β (400 U/mL) or IFN-γ (50 ng/mL) or infected with MRSA (MOI 20) for 24 h subjected to SDS-PAGE and immunoblot analysis of TOM20 with total protein staining achieved by Revert700. **G.** Representative flow plots and gating of MTDR vs. MTG in mock and 24 h MRSA-infected BMDM. **H.** LC-MS acquisition and analysis method overview. **I.** Metabolite peak area plotted as a 10-90 percentile box-whisker plot per sample, showing comparable loading between samples, where the median is indicated with a line and the mean is a plus-sign. **J.** Volcano plot with significantly changed (*P*-value < 0.05) metabolites highlighted as red dots. Gray dots indicate non-significant (*P*-value > 0.05) or low abundance (max peak area < 2000) metabolites. **K.** Significantly changed metabolites in MRSA infection compared to mock (*P*-value < 0.05 and $|\log_2(\text{fold change})| > 0.5$). **L.** Highlighted metabolite peak area quantification across biological replicates. Graphs represent the mean of n ≥ 3 biological replicates with SD error bars. *P*-values were calculated using a paired t-test (**B**), unpaired t-test (**C**, **E**, and **L**). *P < 0.05; **P < 0.01; ***P < 0.001

### Fig S2

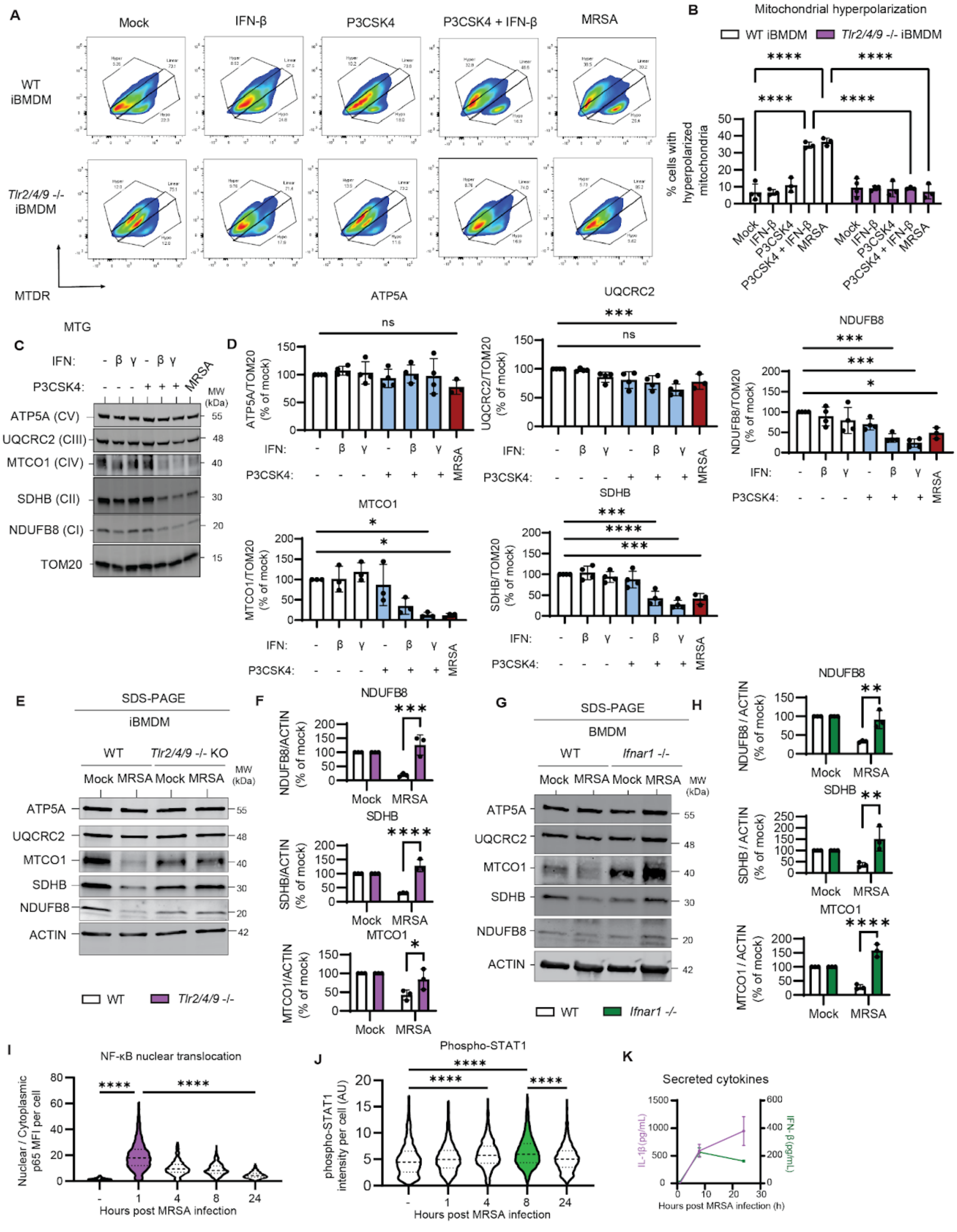
A
Mock
IFN-β
P3CSK4
P3CSK4 + IFN-β
MRSA
WT iBMDM
Tlr2/4/9 -/- iBMDM
MTDR
MTG
B
Mitochondrial hyperpolarization
WT iBMDM
Tlr2/4/9 -/- iBMDM
% cells with hyperpolarized mitochondria
****
Mock
IFN-β
P3CSK4
P3CSK4 + IFN-β
MRSA
C
IFN:
P3CSK4:
MRSA
MW (kDa)
ATP5A (CV)
UQCRC2 (CIII)
MTCO1 (CIV)
SDHB (CII)
NDUFB8 (CI)
TOM20
D
ATP5A
ns
ATP5A/TOM20 (% of mock)
UQCRC2
***
UQCRC2/TOM20 (% of mock)
NDUFB8
NDUFB8/TOM20 (% of mock)
MTCO1
MTCO1/TOM20 (% of mock)
SDHB
****
SDHB/TOM20 (% of mock)
E
SDS-PAGE
iBMDM
WT
Tlr2/4/9 -/- KO
Mock MRSA Mock MRSA
ATP5A
UQCRC2
MTCO1
SDHB
NDUFB8
ACTIN
F
NDUFB8/ACTIN (% of mock)
SDHB/ACTIN (% of mock)
MTCO1/ACTIN (% of mock)
Mock
MRSA
G
BMDM
Ifnar1 -/-
H
NDUFB8 / ACTIN (% of mock)
SDHB / ACTIN (% of mock)
MTCO1 / ACTIN (% of mock)
I
NF-κB nuclear translocation
Nuclear / Cytoplasmic p65 MFI per cell
Hours post MRSA infection
J
Phospho-STAT1
phospho-STAT1 intensity per cell (AU)
K
Secreted cytokines
IL-1β (pg/mL)
IFN-β (pg/mL)
Hours post MRSA infection (h)

**Fig S2. Supplemental data supporting Figure 2. A.** Representative flow plots and gating of MTDR vs. MTG from WT and *Tlr2/4/9* -/- iBMDM treated with TLR2 agonist P3CSK4 with or IFN-β or infected with MRSA (MOI 20) for 24 h. **B.** Percent of cells with hyperpolarized mitochondria per condition per experiment. **C.** Whole cell extracts from BMDM treated with TLR2 agonist P3CSK4 with or without IFN-β or IFN-γ or infected with MRSA (MOI 20) for 24 h subjected to SDS-PAGE and immunoblot analysis of ATP5A, UQCRC2, MTCO1, SDHB, NDUFB8, and TOM20. **D.** Quantification of each respiratory complex subunit relative to TOM20 per condition per experiment as a percentage of the paired mock condition. **E.** Whole cell extracts from WT and *Tlr2/4/9* -/- iBMDM infected with MRSA (MOI 20) for 24 h subjected to SDS-PAGE and immunoblot analysis of ATP5A, UQCRC2, MTCO1, SDHB, NDUFB8, and ACTIN. **F.** Quantification of each respiratory complex subunit relative to ACTIN per condition per experiment for **Fig S2E** as a percentage of the paired mock condition for each cell line. **G.** Whole cell extracts from WT and *Ifnar1* -/- BMDM infected with MRSA (MOI 20) for 24 h subjected to SDS-PAGE and immunoblot analysis of ATP5A, UQCRC2, MTCO1, SDHB, NDUFB8, and ACTIN. **H.** Quantification of each respiratory complex subunit relative to ACTIN per condition per experiment for **Fig S2G** as a percentage of the paired mock condition for each cell line. Violin plots of cell-level quantification of the nuclear:cytoplasmic ratio of NF-κB p65 (**I**) and intensity of phospho-STAT1 (**J**) per cell in CellProfiler, corresponding to the graph in **Fig 2K**. Graphs represent BMDM pooled across three biological replicates comprising analysis of >492 cells with dotted lines indicating the median and quartiles. **K.** ELISA analysis of IL-1β and IFN-β from supernatants of mock, 8 h, or 24 h MRSA infection of BMDM. Graphs represent the mean of $n \geq 3$ biological replicates with SD error bars. *P*-values were calculated using a two-way ANOVA with Sidak's post test (**B**, **F**, and **H**) or a one-way ANOVA (**D**, **I**, and **J**) with Tukey's post test *$P < 0.05$; **$P < 0.01$; ***$P < 0.001$; ****$P < 0.0001$

### Fig S3

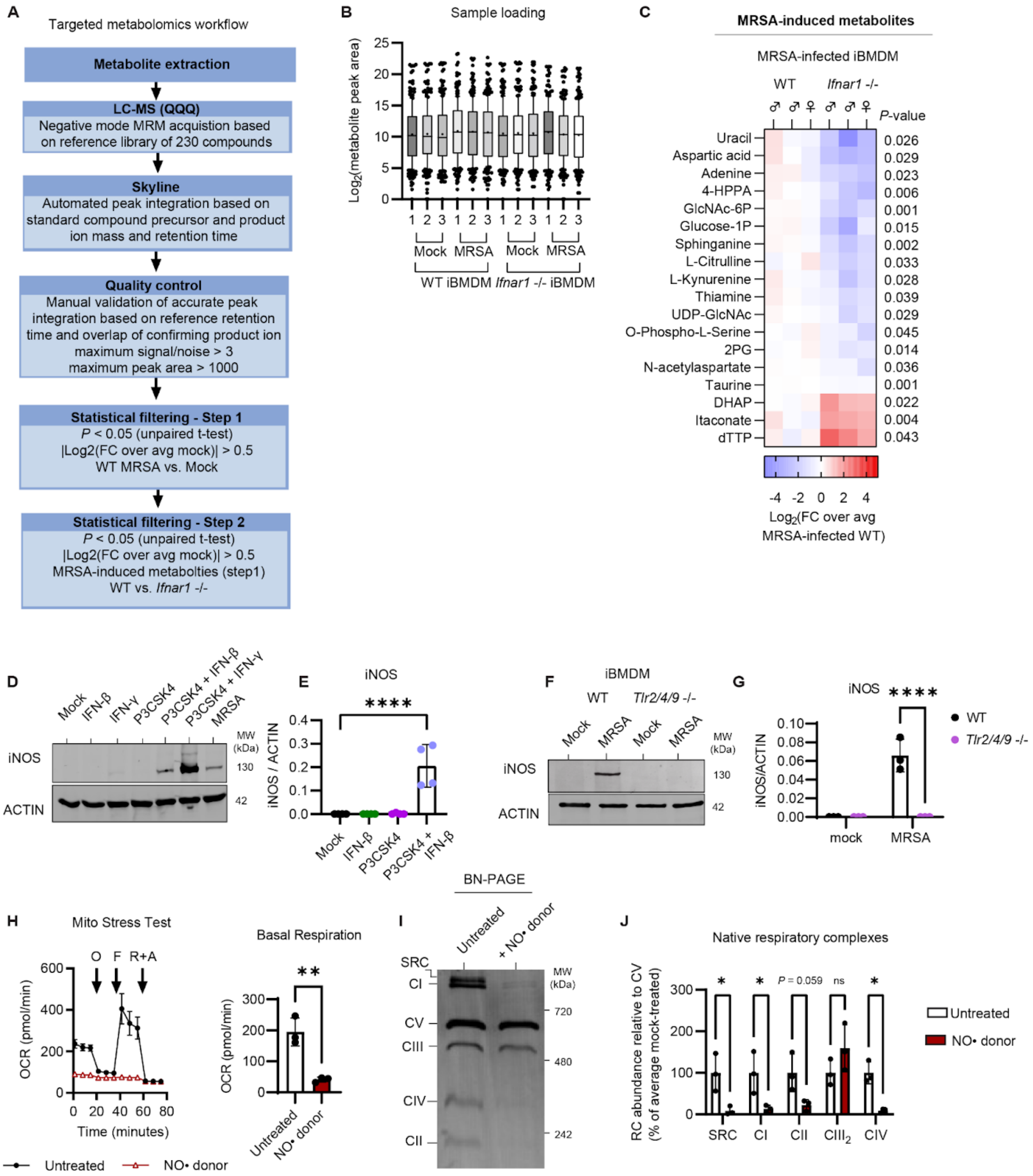
A
Targeted metabolomics workflow
Metabolite extraction
LC-MS (QQQ)
Negative mode MRM acquistion based on reference library of 230 compounds
Skyline
Automated peak integration based on standard compound precursor and product ion mass and retention time
Quality control
Manual validation of accurate peak integration based on reference retention time and overlap of confirming product ion
maximum signal/noise > 3
maximum peak area > 1000
Statistical filtering - Step 1
P < 0.05 (unpaired t-test)
|Log2(FC over avg mock)| > 0.5
WT MRSA vs. Mock
Statistical filtering - Step 2
P < 0.05 (unpaired t-test)
|Log2(FC over avg mock)| > 0.5
MRSA-induced metabolties (step1)
WT vs. Ifnar1 -/-
B
Sample loading
Log2(metabolite peak area)
Mock
MRSA
WT iBMDM
Ifnar1 -/- iBMDM
C
MRSA-induced metabolites
MRSA-infected iBMDM
WT
Ifnar1 -/-
P-value
Uracil 0.026
Aspartic acid 0.029
Adenine 0.023
4-HPPA 0.006
GlcNAc-6P 0.001
Glucose-1P 0.015
Sphinganine 0.002
L-Citrulline 0.033
L-Kynurenine 0.028
Thiamine 0.039
UDP-GlcNAc 0.029
O-Phospho-L-Serine 0.045
2PG 0.014
N-acetylaspartate 0.036
Taurine 0.001
DHAP 0.022
Itaconate 0.004
dTTP 0.043
Log2(FC over avg MRSA-infected WT)
D
Mock
IFN-β
IFN-γ
P3CSK4
P3CSK4 + IFN-β
P3CSK4 + IFN-γ
MRSA
iNOS
ACTIN
MW (kDa)
130
42
E
iNOS
iNOS / ACTIN
****
Mock
IFN-β
P3CSK4
P3CSK4 + IFN-β
F
iBMDM
WT
Tlr2/4/9 -/-
Mock
MRSA
iNOS
ACTIN
MW (kDa)
130
42
G
iNOS
****
iNOS/ACTIN
WT
Tlr2/4/9 -/-
mock
MRSA
H
Mito Stress Test
O
F
R+A
OCR (pmol/min)
Time (minutes)
Untreated
NO• donor
Basal Respiration
OCR (pmol/min)
**
Untreated
NO• donor
I
BN-PAGE
Untreated
+ NO• donor
SRC
CI
CV
CIII
CIV
CII
MW (kDa)
720
480
242
J
Native respiratory complexes
RC abundance relative to CV (% of average mock-treated)
P = 0.059
ns
SRC
CI
CII
CIII2
CIV
Untreated
NO• donor

**Fig S3. Supplemental data supporting Figure 3.** LC-MS-based targeted metabolomics analysis of 230 metabolites in mock or 24 h MRSA-infected (MOI 20) WT or *Ifnar1* -/- iBMDM analyzed in Skyline MS analysis software. **A.** LC-MS acquisition and analysis method overview. **B.** Metabolite peak area plotted as a 10-90 percentile box-whisker plot per sample, showing comparable loading between samples, where the median is indicated with a line and the mean is a plus-sign. **C.** Heat map highlighting significantly different MRSA-induced metabolites in WT vs. *Ifnar1* -/- iBMDM. *P*-value calculated as described in the workflow. **D.** Whole cell extracts from BMDM treated with TLR2 agonist P3CSK4 with or without IFN-β or IFN-γ or infected with MRSA (MOI 20) for 24 h subjected to SDS-PAGE and immunoblot analysis of iNOS and ACTIN. **E.** Quantification of iNOS relative to ACTIN per condition per experiment. **F.** Whole cell extracts from WT and *Tlr2/4/9* -/- iBMDM infected with MRSA (MOI 20) for 24 h subjected to SDS-PAGE and immunoblot analysis of iNOS and ACTIN. **G.** Quantification of iNOS relative to ACTIN per condition per experiment. **H.** Seahorse Mito Stress Test analysis of mock or 24 h nitric oxide (NO•) donor DETA-NONOate-treated BMDM and extracted measurement of basal respiration. **I.** BN-PAGE and immunoblot analysis of native respiratory complexes (RC) from mock or 24 NO• donor-treated BMDM. **J.** Quantification of each native RC relative to Complex V (CV) which does not change in abundance in response to NO•. Graphs represent the mean of $n \geq 3$ biological replicates with SD error bars. *P*-values were calculated using an unpaired t-test (**H**), a one-way ANOVA with Tukey's post test (**E**), or a two-way ANOVA with Sidak's post test (**G** and **J**). *P < 0.05; **P < 0.01; ****P < 0.0001.

### Fig S4

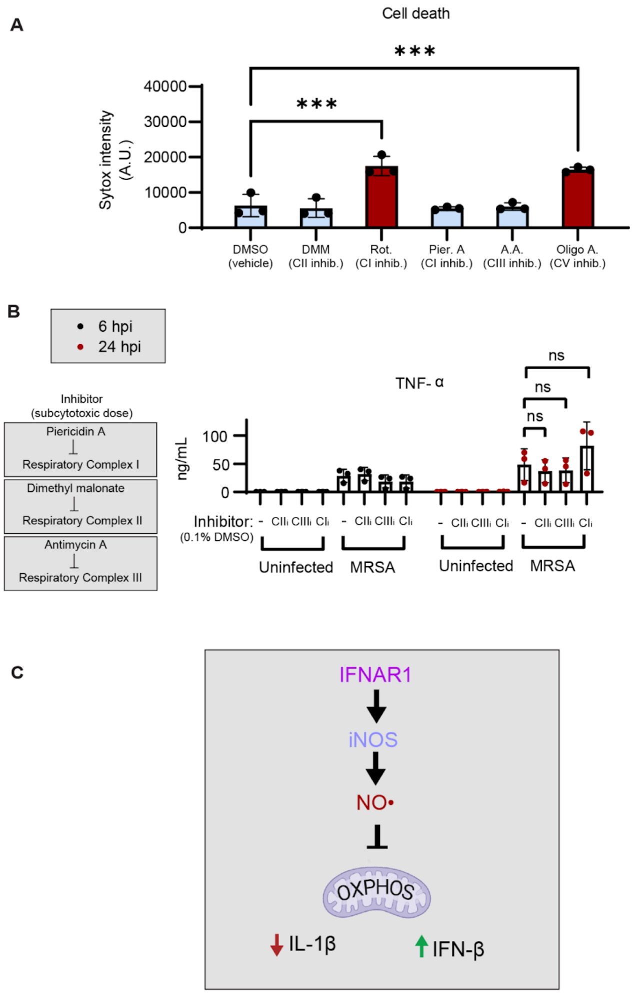
A
Cell death
***
***
Sytox intensity (A.U.)
40000
30000
20000
10000
0
DMSO (vehicle)
DMM (CII inhib.)
Rot. (CI inhib.)
Pier. A (CI inhib.)
A.A. (CIII inhib.)
Oligo A. (CV inhib.)
B
6 hpi
24 hpi
Inhibitor (subcytotoxic dose)
Piericidin A
Respiratory Complex I
Dimethyl malonate
Respiratory Complex II
Antimycin A
Respiratory Complex III
TNF-α
ns
ns
ns
ng/mL
100
50
0
Inhibitor: (0.1% DMSO)
- CIIi CIIIi CIi
- CIIi CIIIi CIi
- CIIi CIIIi CIi
- CIIi CIIIi CIi
Uninfected
MRSA
Uninfected
MRSA
C
IFNAR1
iNOS
NO•
OXPHOS
IL-1β
IFN-β

**Fig S4. Supplemental data supporting Figure 4. A.** Sytox viability assay following BMDM treatment with vehicle control (DMSO), complex I inhibitors rotenone (1 μM) or Piericidin A (100 nM), Complex II inhibitor dimethyl malonate (10 mM), Complex III inhibitor Antimycin A (1 μM), or Complex V inhibitor Oligomycin A (1 μM). Inhibitors color coded in light blue were selected for ELISA analysis due to low cytotoxicity compared to the vehicle control. **B.** ELISA analysis of secreted TNF-α after 6h or 24 h MRSA infection with or without respiratory complex inhibitors, dimethyl malonate (Complex II inhibitor; 10 mM), Antimycin A (Complex III inhibitor; 1 μM), and Piericidin A (Complex I inhibitor; 100 nM). **C.** Model illustrating the pathway of nitric oxide (NO•) generation in MRSA-infected macrophages and the effect of NO• on macrophage metabolism and cytokine responses, created with Biorender.com. Graphs represent the mean of n ≥ 3 biological replicates with SD error bars. *P*-values were calculated using a one-way ANOVA with Tukey's post test (**A**) or two-way ANOVA with Sidak's post test (**B**). No significant differences in TNF-α secretion were identified between MRSA-infected samples at 6 or 24 h post infection. ***P < 0.001

92 **Fig S5**

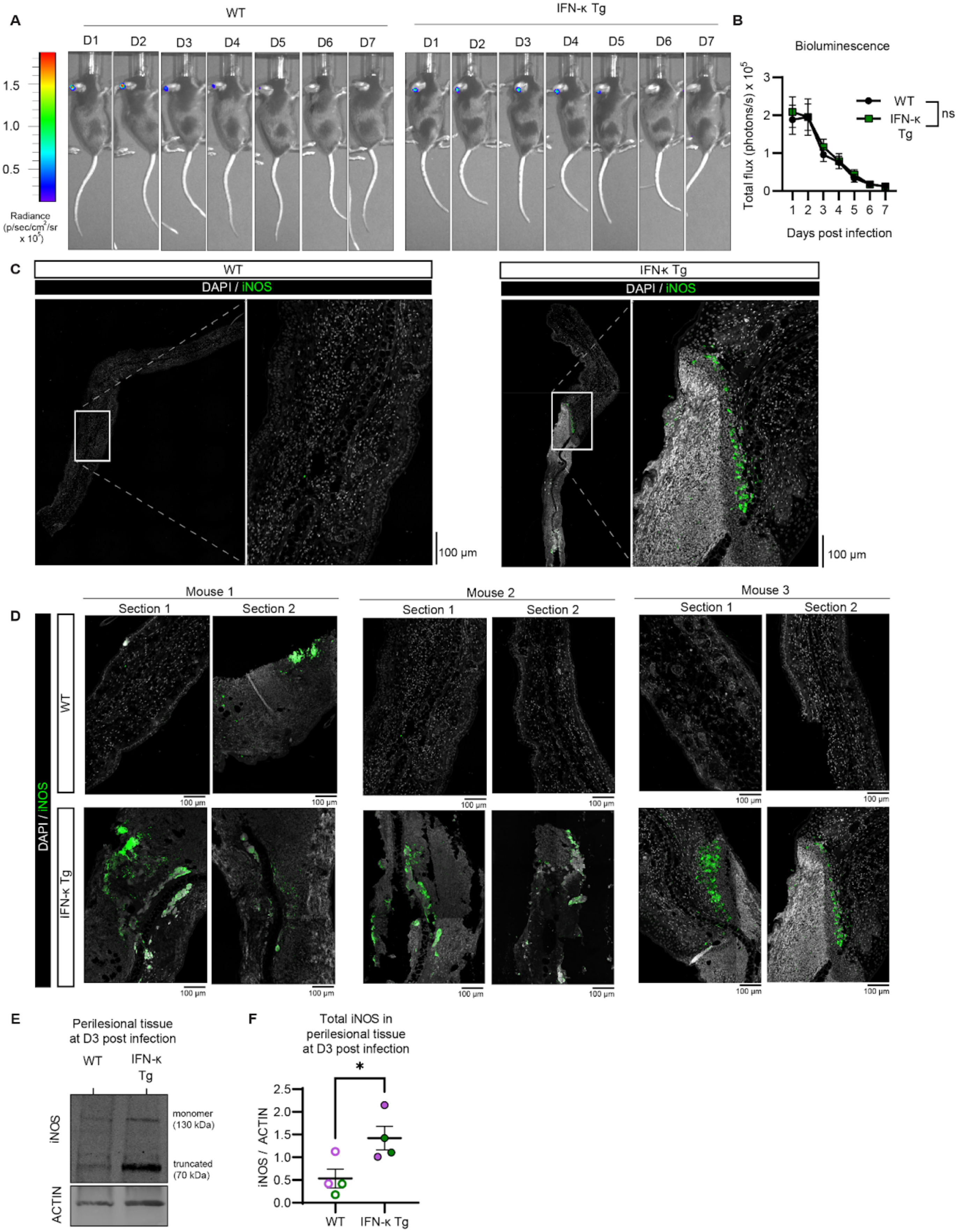

A
WT
IFN-κ Tg
D1
D2
D3
D4
D5
D6
D7
1.5
1.0
0.5
Radiance (p/sec/cm²/sr x 10⁵)
B
Bioluminescence
Total flux (photons/s) x 10⁵
Days post infection
WT
IFN-κ Tg
ns
C
WT
IFNκ Tg
DAPI / iNOS
100 µm
D
Mouse 1
Mouse 2
Mouse 3
Section 1
Section 2
WT
IFN-κ Tg
DAPI / iNOS
100 µm
E
Perilesional tissue at D3 post infection
WT
IFN-κ Tg
iNOS
ACTIN
monomer (130 kDa)
truncated (70 kDa)
F
Total iNOS in perilesional tissue at D3 post infection
iNOS / ACTIN
*
WT
IFN-κ Tg


93

**Fig S5. Supplemental data supporting Figure 6. A.** WT and IFN-κ Tg mice were infected cutaneously with an allergy needle on one ear with bioluminescent MRSA (8e8 CFU; USA300 Lux) and bioluminescence was monitored every 24 h for 7 days with an In Vivo Imaging System Spectrum (IVIS Spectrum). **A.** Representative photograph and bioluminescence images each day for 7 days (D1-7) of WT and IFN-κ Tg mice. **B.** Quantification of bioluminescence radiance as total photons over a fixed region of interest centered around the infected ear. **C.** Immunohistofluorescence visualization of iNOS with DAPI counterstain from automated full section confocal imaging with highlighted zoomed region of day 3 MRSA-infected perilesional (PL) skin, as shown in **Fig 6H**. **D.** Zoomed representative regions of micrographs from multiple immunohistofluorescence stained sections from WT and IFN-κ Tg MRSA PL skin derived from different mice. **E.** Protein extract from MRSA-infected wound and perilesional (PL) skin from WT and IFN-κ Tg mice was subjected to SDS-PAGE and immunoblot analysis of iNOS and ACTIN. Monomeric (130 kDa) and truncated (70 kDa) iNOS species can be detected. **F**. Quantification of total iNOS relative to ACTIN in wound and PL skin. Bioluminescence graphs represent the mean of $n \geq 15$ mice per genotype with SEM error bars. Other graphs represent the mean of $n \geq 3$ mice per genotype with SEM error bars. *P*-values were calculated using a two-way ANOVA with Sidak's post test (**B**) and unpaired t-test (**F**). $^{*}P < 0.05$

### Fig S6

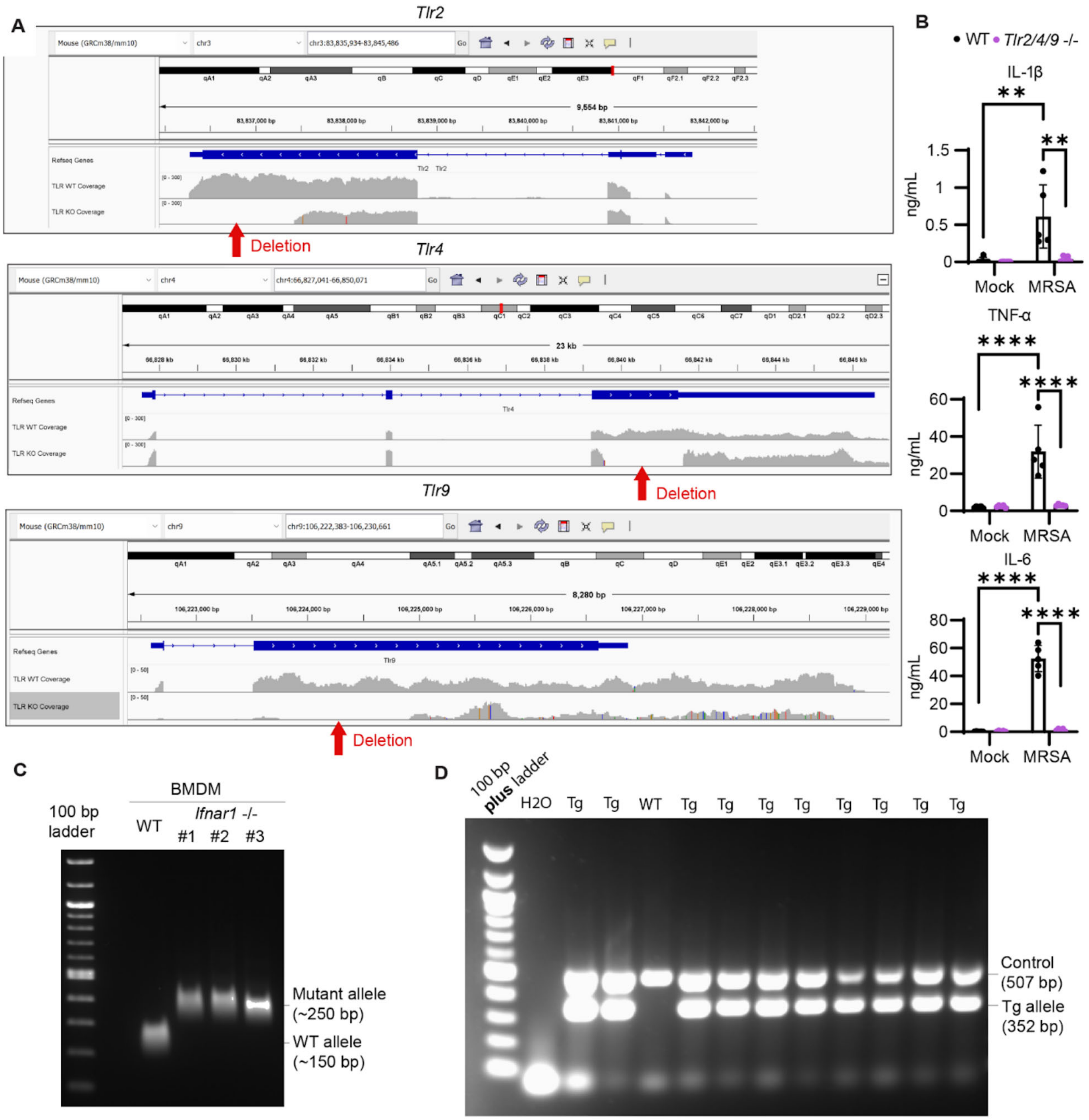
A
Tlr2
Deletion
Tlr4
Deletion
Tlr9
Deletion
B
WT Tlr2/4/9 -/-
IL-1β
ng/mL
Mock
MRSA
TNF-α
IL-6
C
100 bp ladder
BMDM
WT
Ifnar1 -/-
#1
#2
#3
Mutant allele (~250 bp)
WT allele (~150 bp)
D
100 bp plus ladder
H2O
Tg
WT
Control (507 bp)
Tg allele (352 bp)

**Fig S6. Validation of *Tlr2/4/9* -/- macrophages, *Ifnar1* -/- macrophages, and IFN-κ Tg mice. A.** *Tlr2*, *Tlr4*, and *Tlr9* transcript alignment from RNAseq analysis viewed in Integrated Genomics Viewer (IGV), indicating expected exon deletions. **B.** ELISA analysis of IL-1β, TNF-α, and IL-6 from 24 h MRSA-infected WT and *Tlr2/4/9* -/- iBMDM supernatants. **C.** Representative SYBR-stained agarose gel run with PCR-based genotyping results from WT and *Ifnar1*-/- iBMDM. **D.** Representative SYBR-stained agarose gel run with PCR-based genotyping results from tail clips from WT and IFN-κ Tg mice. Band at 507 base pairs (bp) indicates the internal control (present in WT and Tg mice) and the band at 352 bp corresponds to the transgene. Graphs represent the mean of $n \geq 3$ experimental replicates with SD error bars. *P*-values were calculated using a two-way ANOVA with Sidak's post test (**B**). $^{**}P < 0.01$; $^{****}P < 0.0001$
